# Quantification of *nosZ* genes and transcripts in activated sludge microbiomes with novel group-specific qPCR methods validated with metagenomic analyses

**DOI:** 10.1101/710483

**Authors:** DaeHyun D. Kim, Doyoung Park, Hyun Yoon, Taeho Yun, Min Joon Song, Sukhwan Yoon

**Author notes:** Address correspondence to Sukhwan Yoon. Department of Civil and Environmental Engineering, Georgia Institute of Technology, Atlanta, Georgia, USA. Department of Civil and Environmental Engineering, Cornell University, Ithaca, New York, USA.

## Abstract

Substantial N_2_O emission results from activated sludge nitrogen removal processes. The importance of N_2_O-reducers possessing NosZ-type N_2_O reductases have been recognized as the only N_2_O sink *in situ* key to determination of the net N_2_O emissions; however, reliable quantification methods for *nosZ* genes and transcripts have yet to be developed. Here, *nosZ* genes and transcripts in activated sludge tank microbiomes were analyzed with the group-specific qPCR assays designed *de novo* combining culture-based and computational approach. A sewage sample was enriched in a batch reactor fed continuous stream of N_2_ containing 20-10,000 ppmv N_2_O, where 14 genera of potential N_2_O-reducers were identified. All available amino acid sequences of NosZ affiliated to these taxa were grouped into five subgroups (two clade I and three clade II groups), and primer/probe sets exclusively and comprehensively targeting the subgroups were designed and validated with *in silico* PCR. Four distinct activated sludge samples from three different wastewater treatment plants in Korea were analyzed with the qPCR assays and the results were validated by comparison with the shotgun metagenome analysis results. With the validated qPCR assays, the *nosZ* genes and transcripts of six additional activated sludge samples were analyzed and the results of the analyses clearly indicated the dominance of two clade II *nosZ* subgroups (*Flavobacterium*-like and *Dechloromonas*-like) among both *nosZ* gene and transcript pools.

## Introduction

Nitrous oxide (N_2_O) is one of the three major greenhouse gases with the largest contributions to global warming, along with CO_2_ and CH_4_.^1^ Although the contribution of N_2_O is estimated to be only ∼6% of the net greenhouse gas emissions in terms of CO_2_eq, which is far less than those of CO_2_ and CH_4_, eliminating one molecule of N_2_O from the atmosphere has the same merit as removing ∼300 molecule of CO_2_ due to its high global warming potential. Besides, N_2_O has also been the most consequential ozone depletion agent.^2, 3^ Thus, global efforts to curb the increase in atmospheric N_2_O concentration are necessitated for sustainable future. A better understanding of the biogeochemical reactions functioning as N_2_O sources and sinks in nitrogen-rich anthropogenic environments, e.g., fertilized agricultural soils and wastewater treatment plants (WWTPs), is especially important for N_2_O emission mitigation, as nitrification and denitrification, the biological reactions serving as the major sources of N_2_O, occur at consistent a basis in such environments and thus, they have been estimated as the hotspots of N_2_O emission to the atmosphere.^4, 5^

Particularly, the biological N_2_O reduction mediated by Nos-type nitrous oxide reductases (NosZ) has recently attracted immense scientific attention as the sole sink of N_2_O on the Earth’s surface at non-elevated temperature.^6–9^ This reaction had been, for long, known merely as one of the stepwise reactions constituting the denitrification pathway.^4, 10^ Only recently has the N_2_O-to-N_2_ reduction been recognized as an independent energy-conserving reaction, as diverse organisms possessing either conventional clade I and newly-discovered clade II *nosZ* were found to be capable of growth with N_2_O as the sole electron acceptor.^9, 11–13^ Several independent research groups have reported difference between clade I and clade II *nosZ*-possessing organisms in terms of their affinities to N_2_O, and more specifically, the organisms with *nosZ* closely affiliated to *Dechloromonas* spp. have been reported with particularly low whole-cell Michaelis constants (K_m,app_), suggesting that this group of organisms may be involved in *in situ* consumption of low-concentration N_2_O produced via diverse biotic and abiotic processes ^9, 13–16^. The microbial community developed in the laboratory-scale biofilter treating 100 ppmv N_2_O included the clade II N_2_O reducer *Flavobacterium* spp. as the most abundant *nosZ*-carrying organisms, also supporting the significance of clade II N_2_O as an important N_2_O sink.^17, 18^

The NosZ-mediated N_2_O reduction has recently been recognized to have an immensely important role in various engineered wastewater treatment systems as the sole sink of N_2_O produced from denitrification and nitrification.^12, 19–23^ In these recent studies, attempts have been made to correlate N_2_O reduction activities or net N_2_O emissions to *nosZ* gene/transcript abundances or the *nosZ*-to-(*nirS*+*nirK*) abundance ratios. Most, if not all, of these *nosZ* gene quantifications were performed using SYBR Green quantitative polymerase chain reactions (qPCR) targeting the clade I (1840F: 5’-CGCRACGGCAASAAGGTSMSSGT-3’/ 2090R: 5’-CAKRTGCAKSGCRTGGCAGAA-3’) and clade II *nosZ* (nosZ-II-F: 5’-CTIGGICCIYTKCAYAC-3’ / nosZ-II-R: 5’-SKSACCTTITTRCCITYICG-3’).^24, 25^ The reliability of these *nosZ* quantification results were questionable, due to unverified specificity and low amplification efficiency, and even the simplest questions as to whether clade I or clade II *nosZ* were more abundant in nutrient removal bioreactors remains controversal.^12, 26, 27^ In this study, we have developed TaqMan-based qPCR reactions targeting four groups (two clade I *nosZ* groups and two clade II *nosZ* groups) of *nosZ* with amplification efficiency above 90% and verified their coverage and specificity by comparing the quantification data with the results of shotgun metagenome analyses. Anoxic activated sludge samples from anoxic tanks of six conventional wastewater treatment plants in Korea with A2O (anaerobic-anoxic-oxic) configuration were then analyzed with these novel qPCR assays, and without exception, clade II domination of *nosZ* gene and transcript pools was verified.

### Materials and methods Sample collection

The wastewater inoculum for the N_2_O enrichment experiments were grab-sampled from the anoxic section of the activated sludge tank at Daejeon municipal WWTP (36°23’5” N 127°24’28” E) in September 2016 (denoted as Daejeon1). Three activated sludge samples for validation of the qPCR assays by comparison with the metagenomics data were collected at the same WWTP (Daejeon2) in February 2019 and two other activated sludge WWTPs located in Gwangju and Gapyeong (35°09’22.4″N 126°49’51.6″E and 37°49’00.1″N 127°31’13.0″E, respectively) in January and February of 2019, respectively. Activated sludge samples from anoxic tanks of six other A2O (anaerobic/anoxic/oxic) WWTPs were then collected for group-specific quantification of *nosZ* genes and transcripts with the developed qPCR assays (Figure S1). Each wastewater sample for analyses of DNA was collected in a 2-L polyethylene bottle filled up to the brim to minimize oxygen ingress. The samples for quantification of *nosZ* transcripts were immediately mixed with the same volume of methanol for RNA fixation.^28^ The sample bottles were immediately placed in a cooler and transported to the laboratory, where they were stored at -80°C until use.

### Fed-batch enrichment of activated sludge samples and identification of active N_2_O-reducing groups

A simple fed-batch bioreactor was constructed for enrichment of active N_2_O-reducers in the Daejeon1 wastewater sample (Figure S2). A 500-mL glass bottle with a side port (Duran, Mainz, Germany) was fitted with a GL45 cap with three ports. Two of the ports were used as the inlet for N_2_O-carrying gas and the gas outlet from the bottle and the remaining port was used for aqueous phase sampling. The side port of the glass bottle was used for sampling of the gaseous phase. The modified MR-1 medium was prepared by adding per 1 L of deionized water, 0.5 g NaCl, 0.41 g sodium acetate, 0.23 g KH_2_PO_4_, 0.46 g K_2_HPO_4_, 0.026 g NH_4_Cl, 1 ml of 1000X trace metal solution, and 1 mL of 1000X vitamin stock solution.^29^ The reactor with 200-mL medium (40% of the total reactor volume) was continuously supplied with 0 ppmv, 20 ppmv, 200 ppmv, or 10,000 ppmv N_2_O prepared in >99.999% N_2_ gas (Samoh Specialty Gas, Daejeon, South Korea) for 30 hours before inoculation. After inoculating the medium with 2 mL of the Daejeon1 sample, the same gas was bubbled through the medium at the volumetric flowrate of 20 mL min^-1^ to provide the sole electron acceptor N_2_O to the microbial culture and maintain the reactor at anoxic condition. The reactor operated with >99.999% N_2_ gas served as the control to confirm that the microbial consortia were enriched with N_2_O as the electron acceptor and the contribution of O_2_ contamination to microbial growth was kept to minimal. The N_2_O concentration in the headspace of the reactor was monitored with a HP6890 Series gas chromatography fitted with an HP-PLOT/Q column and an electron capture detector (Agilent, Palo Alto, CA), with the injector, oven and detector temperatures set to 200, 85, and 250 °C, respectively.^30^ The O_2_ concentration in the fed-batch reactor was monitored with a FireSting-O_2_ oxygen meter (Pyroscience, Aachen, Germany); however, due to the relatively high detection limit of the sensor (the gas phase concentration of ∼0.1% v/v), complete absence of O_2_ was not guaranteed.

The growth of bacterial population in the reactor was monitored with qPCR using TaqMan chemistry targeting the conserved region of eubacterial 16S rRNA genes. At each sampling time point, an 1.5-mL aliquot was collected from the aqueous phase of the reactor and DNA was extracted from the pellets using DNeasy Blood & Tissue Kit (Qiagen, Hilden, Germany) following the protocol provided by the manufacturer. Quantitative PCR was performed with the 1055f (5’-ATGGCTGTCGTCAGCT-3’) / 1392r (5’-ACGGGCGGTGTGTAC-3’) / Bac1115Probe (5’-CAACGAGCGCAACCC-3’) primer and probe set targeting a conserved region of bacterial 16S rRNA genes using a QuantStudio3 platform (Thermo Fisher Scientific, Waltham, MA).^31^ Incubation was halted when the population of the enrichment reached a plateau, as indicated by three consecutive measurements without increased gene counts. The reactor was dismantled and the aqueous phase was collected for microbial composition analysis. The hypervariable V6−8 region of the 16S rRNA gene was amplified with 926F: 5′-AAACTYAAAKGAATTGRCGG-3′ / 1392R: 5′-ACGGGCGGTGTGTRC-3′ primer set and MiSeq sequencing of the amplicons was outsourced to Macrogen Inc. (Seoul, Korea). The raw sequence reads were deposited in the NCBI short reads archive (SRA) database (accession: PRJNA552413) and were processed using the QIIME pipeline v 1.9.1 (detailed computational method provided in the supplementary information).

### Design of degenerate primers and probes for group-specific qPCR of *nosZ* genes

The genera assigned to the OTUs with the relative abundances higher than 0.3% in any of the reactor microbial communities were selected. All *nosZ* gene sequences and corresponding translated NosZ amino acid sequences belonging to the organisms affiliated to these genera were extracted from the Uniprot (www.uniprot.org) database (accessed in March 2017) (Table S1). The curated pools of *nosZ*/NosZ sequence data included, in total, 174 nucleotide sequences and the corresponding amino acid sequences from 14 distinct genera. Subsequently, a multiple sequence alignment was performed with these amino acid sequences, using MUSCLE algorithm with the parameters set to the default values.^32^ The NosZ phylogenetic tree was constructed using the neighbor-joining method in MEGA 7.0 with the bootstrap value set to 500.^33^ The *nosZ* gene sequences were clustered into five groups (NosZG1-5) according to the positions of the corresponding NosZ sequences in the phylogenetic tree, which consisted of five phylogenetically distinct subbranches.

**Table 1.**
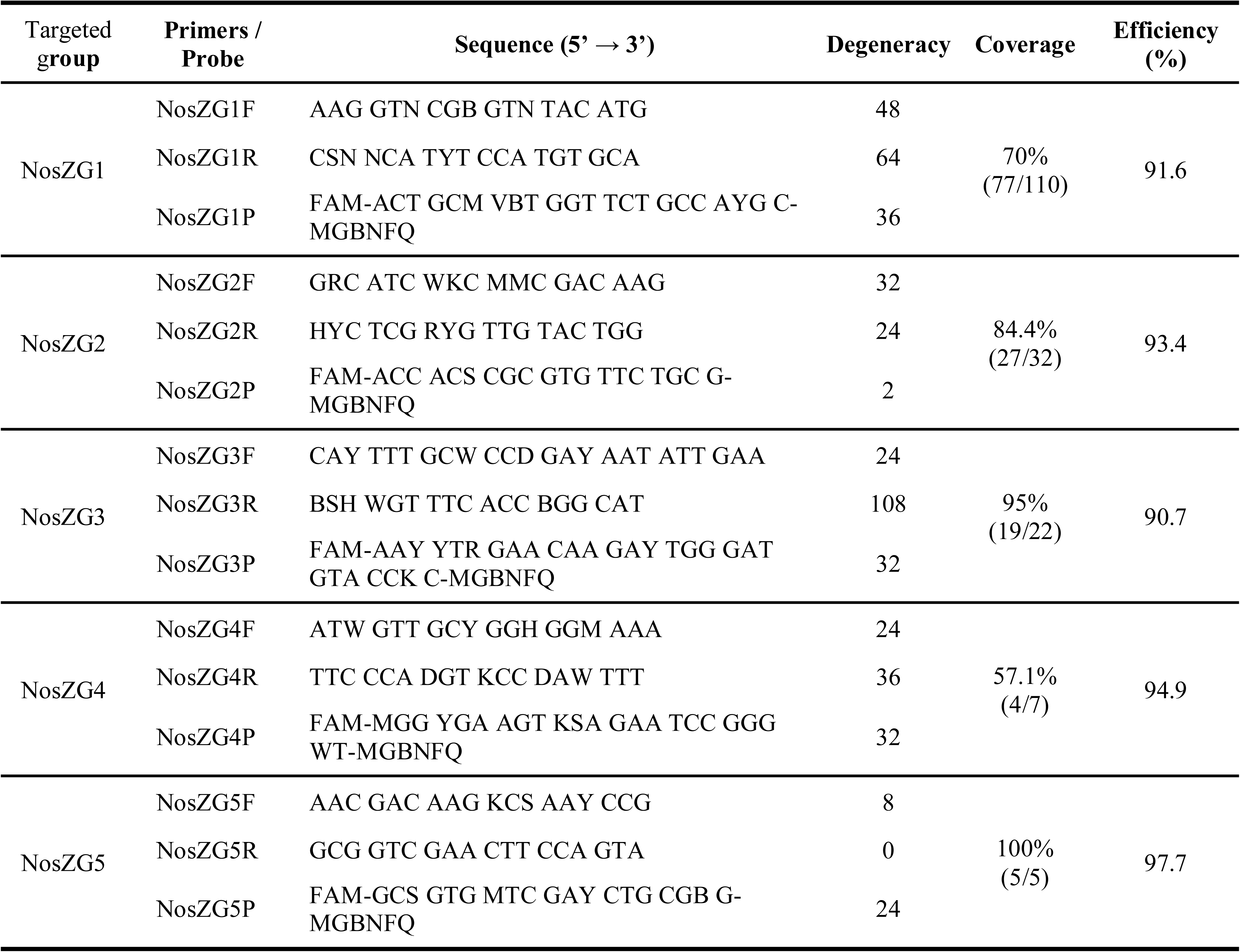
The primers/probe sets developed in this study for group-specific quantification of *nosZ* genes

For each *nosZ* group, a primers and probe set was designed using the PriMux software to comprehensively and exclusively target the *nosZ* gene sequences within the group.^34^ The parameters were modified from the default values to obtain the optimal candidate degenerate oligonucleotide sequences for qPCR (length: 18-24 bp, amplicon size: 80-400 bp, T_m_: 56-62°C for primers and 68 - 72°C for probes). Several candidate primers and probe sets were generated for each *nosZ* group, implementing the ***min***, ***max***, and ***combo*** algorithms of the Primux software. The performance of each candidate primers and probe set was predicted with *in silico* PCR performed with the simulate_PCR software against all complete genome sequences of the organisms belonging to each of the five *nosZ* groups (Table S1).^34, 35^ Coverage within the target *nosZ* group and mutual exclusivity across the groups were the two major criteria for assessment of the candidate primers and probe sets. The *in silico* PCR tests were also performed against the complete genomes of 50 bacterial strains lacking *nosZ* to preclude the possibility of unspecific amplification.

### Construction of calibration curves for the designed primer and probe sets using model N_2_O reducer strains

A model organism from each group of the selected *nosZ* gene(s) was used for construction of the calibration curve. The selected model organisms were *Pseudomonas stutzeri* DCP-Ps1 (NosZG1), *Acidovorax soli* DSM25157 (NosZG2), *Flavobacterium aquatile* LMG4008 (NosZG3), *Ignavibacterium album* JCM16511 (NosZG4), and *Dechloromonas aromatica* RCB (NosZG5). *Acidovorax soli* DSM25157 and *F. aquatile* LMG4008 were acquired from Korean Collection for Type Cultures and *I. album* JCM16511 from Japanese Collection of Microorganisms. The axenic batch cultures of these organisms were prepared as previously described in the literature or using the media and incubation conditions specified by the distributors. The cells were harvested at OD_600nm_ = 1.1 and the *nosZ* gene(s) in the DNA extracted from the cell pellets were amplified using the designed primers. The calibration curve for each qPCR reaction was constructed using ten-fold serial dilutions of PCR^®^ 2.1 vectors (Invitrogen, Carlsbad, CA) carrying the *nosZ* amplicons. A uniform thermocycle was used for the qPCR: 95°C for 10 min and 40 cycles of 95 °C for 30 s, 58 °C for 60 s, and 72 °C for 60 s.

### Group specific quantitative PCR targeting *nosZ* gene and transcripts in activated sludge samples

The group-specific quantification of the *nosZ* genes and transcripts in the activated sludge samples were performed with the designed primer and probe sets (Table 1) on a QuantStudio 3 real-time PCR instrument using TaqMan detection chemistry (FAM as the reporter and NFQ-MGB as the quencher). DNeasy Blood & Tissue Kit (QIAGEN) was used to extract DNA from the activated sludge samples. From the methanol-treated samples, RNA was extracted with RNeasy Mini Kit (Qiagen), purified with DNase I (Qiagen) and RNeasy MinElute Cleanup Kit (Qiagen), and reverse-transcribed with Superscript^®^ III (Thermo Fisher Scientific), as described previously.^36^ The luciferase control mRNA (Promega, Madison, WI, USA) was added as the internal standard to account for RNA loss during the process. Each 20-μL qPCR reaction mix contained 10 μL of 2X TaqMan master mix (Applied Biosystems, Foster city, CA, USA), 5 μM each of the forward and reverse primers, 0.5 μM of the probe, and 2 μL of the DNA or cDNA. Calibration curves prepared with dilution series of the PCR amplicons were used to calculate the copy numbers of the targeted genes from the C_t_ values. Eubacterial 16S rRNA genes in the extracted DNA samples were quantified using the 1055F/1392R/Bac115Probe set for quantification of total bacterial population in the activated sludge samples (Table S2). The copy numbers of the targeted *nosZ* genes in the wastewater samples were normalized with the 16S rRNA gene copy numbers to facilitate comparison with the relative abundances of the *nosZ* groups from the metagenomic analyses.

The PCR amplicons of the Daejeon1 sample amplified with NosZG1-5 primer sets were sequenced using Illumina Miseq platform (San Diego, CA) at the Center for Health Genomics and Informatics at University of Calgary. The raw sequence reads have been deposited in the SRA database (accession number: PRJNA552418). After quality trimming and merging of the paired-end sequences, the sequences without the probe-binding region were removed, and the remaining reads were clustered into OTUs with 0.97 cut-off using cd-hit-est v. 4.6.^37^ The OTUs were annotated using blastx against the bacterial Refseq database downloaded in June, 2018, with the e value cut off set to 10^-3^ and word size to 3 and no seg option selected,

### Computational quantification of *nosZ* genes from shotgun metagenomes of the activated sludge samples

The DNA samples for shotgun metagenome sequencing were extracted with DNeasy PowerSoil Kit (Qiagen) from 50 mL each of the four activated sludge samples collected for validation of the qPCR assays. Sequencing of the metagenomic DNA was performed at Macrogen Inc., where Hiseq X Ten sequencing platform (Illumina, San Diego, CA) was used for generating 5-10 Gb of paired-end reads data with 150-bp read length. The raw sequence reads have been deposited in the NCBI short reads archive (SRA) database (accession numbers: PRJNA552406).

The raw reads were then processed using Trimmomatic v0.36 software with the parameters set to the default values.^38^ The trimmed reads were translated *in silico* into amino acid sequences using all six possible reading frames and screened for clade I and II *nosZ* sequences using hidden Markov models (HMM). The two HMM algorithms for clade II *nosZ* were downloaded from the Fungene database (accessed in October 2016). As the HMM for clade I *nosZ* in the database had specificity issues, the HMM algorithm was constructed *de novo* using *hmmbuild* command of HMMER v3.1b1. The clade I NosZ sequences used to build the HMM were manually curated from the pool of NosZ sequences downloaded from the NCBI database (accessed in October 2016), to represent diverse subgroups within the clade (Table S3).^8, 11^ The candidate partial NosZ sequences were extracted from the translated shotgun metagenome reads using the *hmmsearch* command of HMMER v3.1b1 with the e value cutoff set to 10^-5^. The nucleotide sequence reads corresponding to the extracted partial NosZ sequences (in separate bins for clade I and clade II NosZ) were assembled into contigs using metaSPAdes v3.12.0 with parameters set to default values.^39^ The assembled contigs with lengths shorter than 200 bp were filtered out. The overlapping contigs appearing in both clade I and clade II NosZ bins were identified by clustering the two sets of contigs against each other with a nucleotide identity cutoff of 1.0. These overlapping contigs were manually called to the correct bin according to the BLASTX results. The trimmed sequence reads were then mapped onto the contigs in the *nosZ* bins using Bowtie2 v2.2.6, yielding sequence alignments for the contigs and the mapped reads. The alignment files were further processed using samtools v0.1.19, and the PCR duplicates were removed using MarkDuplicates function of picard-tools v1.105.^40^ The sequence coverages of the contigs were calculated using bedtools v2.17.0. The number of reads mapped on to the contigs were normalized with the lengths of the respective contigs. The contigs were assigned taxonomic classification using blastx, and distributed to the NosZG1-5 bins based on their taxonomic affiliations (Table S4). The contigs without matching sequence were binned as ‘other *nosZ* sequences’. The attempt to use 16S rRNA sequences extracted with Meta-RNA and assembled with EMIRGE as the template for mapping was not successful, as the coverage of the extracted 16S rRNA sequences turned out to be an order of magnitude lower than the *rpoB* coverage (Data not shown).^41, 42^ Thus, the sequence coverage of *rpoB* gene, a single-copy housekeeping gene, was used for normalization of the *nosZ* abundance data. The HMM algorithm for *rpoB* was downloaded from the Fungene database.

## Results

### Active N_2_O reducers enriched in fed-batch incubation with varying N_2_O concentrations

The microbial communities of the three N_2_O-reducing enrichments, each prepared with different N_2_O concentrations, were analyzed to identify active N_2_O-reducing groups of microorganisms (Table S5). Screening for operational taxonomic units (OTUs) with >0.3% abundance in any of the three enrichments yielded 69 OTUs assigned to 33 genera in total, and 14 of these genera were identified with phylogenetic subgroups harboring clade I or clade II *nosZ*. The OTUs belonging to these putatively *nosZ*-harboring genera amounted to 50.8% – 63.2% of the total microbial population in the enrichments. The abundances of the genera putatively harboring clade II *nosZ* were observed to be greater than those of the genera harboring clade I of *nosZ* (Table S5). In the enrichment incubated with 20 ppmv N_2_O, *Cloacibacterium* (18.9%), *Flavobacterium* (14.2%), and *Acidovorax* (13.5%) were identified as the dominant genera. *Dechloromonas* (17.3%) and *Flavobacterium* (15.8%) were the dominant genera in the 200 ppmv enrichment, and *Dechloromonas* was the predominant population in the enrichment incubated with 10,000 ppmv of N_2_O, constituting 46.0% of the total microbial population.

### Designing of the degenerate *nosZ* primer/probe sets

With the 174 translated NosZ sequences affiliated to the 14 genera putatively harboring active N_2_O-reducing organisms identified in the activated sludge enrichments, a phylogenetic tree composed of five distinct branches (NosZG1 – NosZG5) was constructed (Figure S3). NosZG1 and NosZG2 were identified as clade I NosZ and NosZG3-NosZG5 as clade II NosZ. After the iterative process of designing candidate primers and probe sets and performing *in silico* PCR tests, the final sets of degenerate primers/probe sets were designed to comprehensively and exclusively target the five *nosZ* groups (Table 1). The *in-silico* PCR performed against 174 genomes from which the target *nosZ* sequences were extracted and 50 genomes without *nosZ* confirmed the high levels of coverage (57.3-100%) and complete exclusivity for all five degenerate primers/probe sets (Table 1). The qPCR calibration curves were constructed with the selected model organisms (Figure S4, Table S6), and despite the high levels of degeneracy (up to 110592), amplification efficiencies above 90% were attained for all five primer and probe sets after rigorous optimization process.

### Cross-checking of the group-specific qPCR quantification of *nosZ* genes with shotgun metagenome analyses

The *nosZ* gene abundances in four distinct activated sludge samples were quantified using the newly designed qPCR assays, and the *nosZ* sequences extracted from the shotgun metagenomes of these same samples were quantitatively analyzed in parallel, for cross-checking of the qPCR results (Figure 1). The qPCR results of the four activated sludge samples invariably showed the dominance of the clade II *nosZ* genes (NosZG3 and NosZG5) over clade I *nosZ* genes (NosZG1 and NosZG2), with at least five-fold higher copy numbers (Figure 2). The qPCR targeting the NosZG4 group failed to amplify the clade II *nosZ* belonging to this group. The NosZG4 primers/probe set was designed from a single complete genome (*Ignavibacterium album*) and six sequences from metagenome assembled genomes (MAGs), which may have been insufficient to cover the sequence divergence of this *nosZ* subgroup.

**Figure 1.**
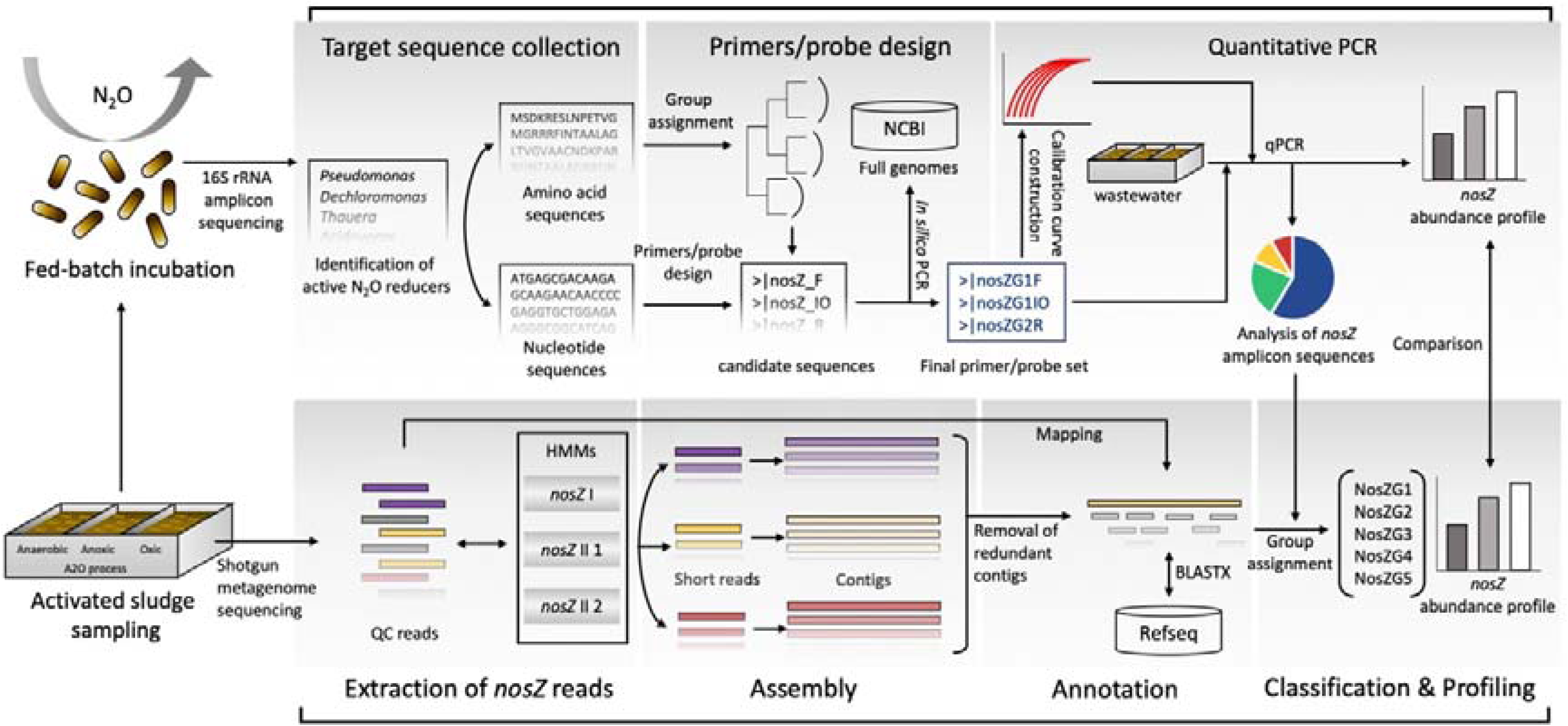
Flow chart for designing of the group-specific *nosZ* primer/probe sets and cross-validation with *nosZ* sequence data extracted from shotgun metagenome data

**Figure 2.**
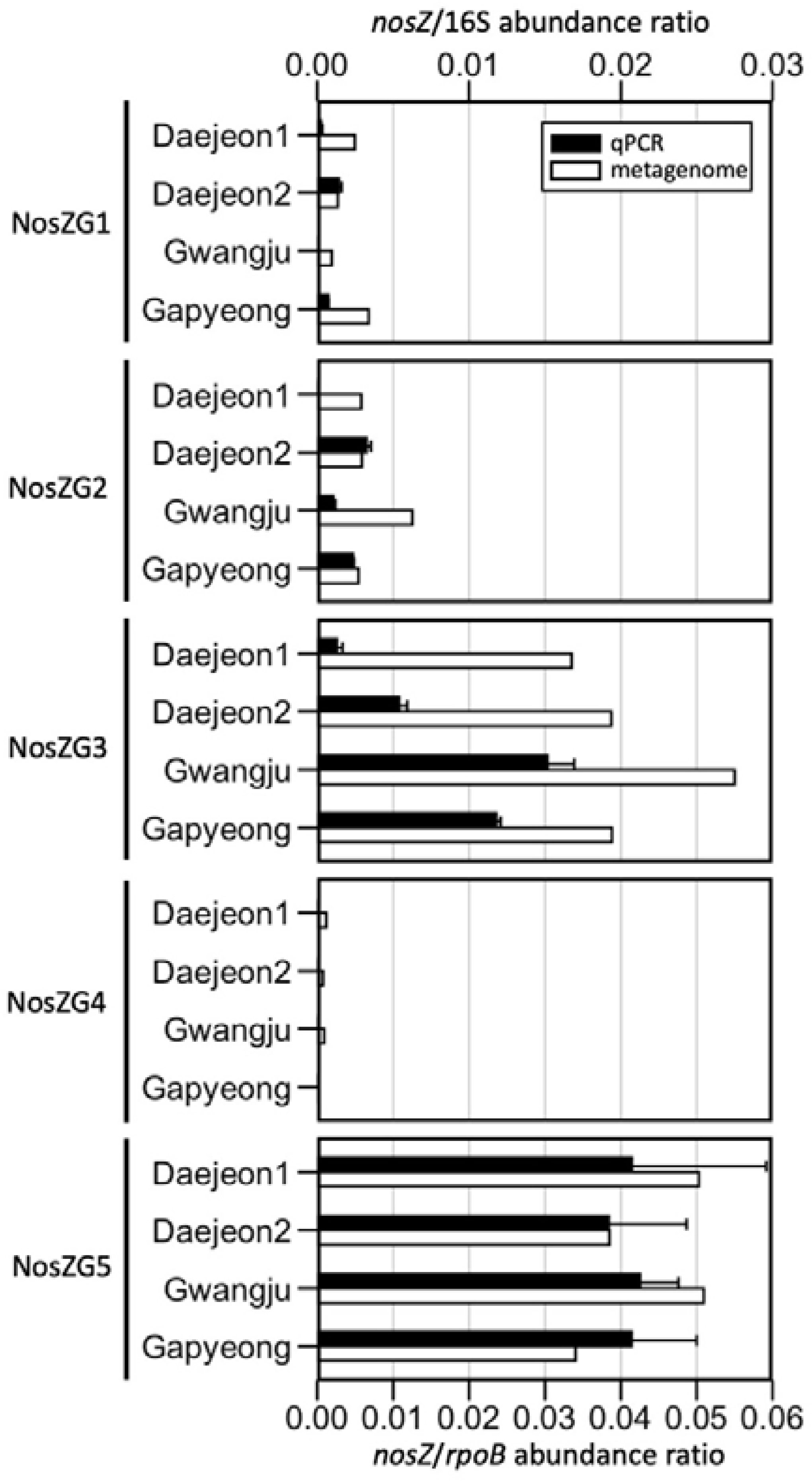
The relative abundances of NosZG1-5 *nosZ* genes as quantified using qPCR and metagenome analysis. The *nosZ* copy numbers from the qPCR assays were normalized with the copy numbers of eubacterial 16S rRNA genes. The coverages of *nosZ* genes in the metagenome data were normalized with the coverages of *rpoB* genes. The presented qPCR quantification data are the averages of triplicate samples processed seperately through extraction and qPCR procedures, with the error bars representing their standard deviations.

The distribution of the *nosZ* sequences extracted from the shotgun metagenomes (Table S7) were also severely biased towards clade II. The compositions of the *nosZ* genes exhibited high level of similarity across the activated sludge samples. The calculated relative abundances of the two most abundant groups, NosZG3 and NosZG5, were relatively consistent across the samples, varying by less than 1.7-fold. The contributions of NosZG1 and NosZG2 to the total *nosZ* gene abundances were minor in all of the samples and largely variable. The fold differences between the samples with the highest relative abundance of NosZG1 and NosZG2 and those with the lowest abundance were 3.5 and 2.3, respectively. The *nosZ* genes sorted as NosZG4 constituted <1.2% of the total *nosZ* genes grouped as NosZG1-NosZG5, suggesting that this *nosZ* group has less significant role in N_2_O reduction in activated sludge tanks than other analyzed groups.

As the qPCR data were normalized with the eubacterial 16S rRNA copy numbers and the metagenome-derived relative abundance data were normalized with the sequence coverages of the single-copy housekeeping gene *rpoB*, direct comparison of the outcomes was not possible. Theoretically, however, the ratios of the *nosZ*/*rpoB* (metagenome) to *nosZ/*16S (qPCR) could be used as indicators of reliability of the qPCR assays, with consistency in the ratios across the *nosZ* groups indicating high reliability. The *nosZ*/*rpoB-*to-*nosZ/*16S ratios were within a narrow range between 1.6 and 2.4 for NosZG5 across the four activated sludge samples. For an unidentified reason, broad gaps between the NosZG1-3 qPCR and the metagenomics data were observed with the Daejeon1 sample. Excluding these results, the *nosZ*/*rpoB*-to-*nosZ*/16S ratio varied from 3.3 to 7.2 for NosZG3, and the *nosZ*/*rpoB*-to-*nosZ*/16S ratios of NosZG1 and NosZG2 ranged from 1.9 to 2.1 and from 1.9 to 9.2, respectively. The *nosZ*/16S-to-*nosZ*/*rpoB* ratios of 12 out of the 16 qPCR assays performed (excluding NosZG4) were within the range between 1.5 and 9.2, indicating that the group-specific qPCR assays enabled reliable quantification of *nosZ* genes.

The *nosZ* amplicons of the Daejeon1 sample, amplified with the newly-designed qPCR primers, were sequenced and analyzed to check whether the qPCR reactions were comprehensive and mutually exclusive (Figure 3). As the NosZG4 primers failed to amplify the targeted genes, NosZG4 was excluded from the downstream analysis. The *nosZ* genes of putative active N_2_O reducers captured by the NosZG1 and NosZG2 primer sets accounted for 54.2% of the entire pool of the clade I *nosZ* genes extracted from the shotgun metagenome. The NosZG3 and NosZG5 primer sets captured 63.4% of the clade II *nosZ* genes recovered from the metagenome (Figure S5). No amplification outside the targeted group occurred for any of the primers, confirming the complete mutual exclusivity of these primers. Each of the qPCR assays were comprehensive within its target groups, as analyzed *nosZ* amplicon sequences covered all genera originally targeted by each primer and probe set.

**Figure 3.**
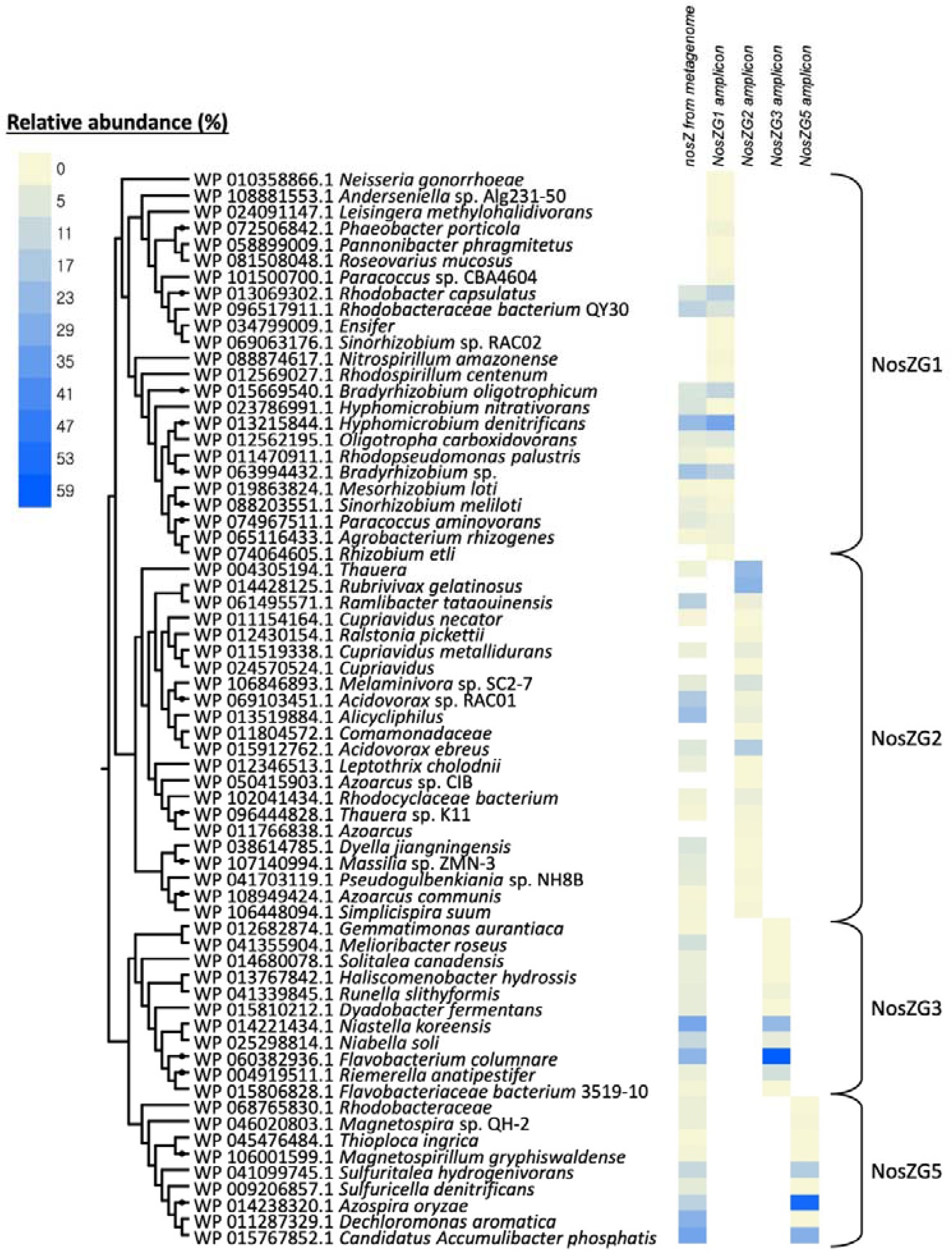
The phylogenetic tree constructed with the reference *nosZ* sequences of the taxa matching OTUs generated from the partial *nosZ* sequences amplified with NosZG1-5 primers. The phylogenetic tree was generated using neighbor-joining method with 500 bootstrap replications. The heatmap shows the relative abundances of the OTUs within each *nosZ* group, computed from the amplicon sequencing data and the *nosZ* sequence data extracted from the metagenome

Overall, the OTUs identified to be abundant from sequence analyses of the PCR amplicons coincided with the dominant *nosZ* OTUs from shotgun metagenome analysis. The OTUs affiliated to *Rhodobacter capsulatus*, Rhodobacteraceae bacterium QY30, *Hyphomicrobium denitrificans*, and *Bradyrhizobium* sp. were the dominant OTUs among the OTUs NosZG1 amplicons. The two most abundant *nosZ* OTUs among the NosZG3 amplicons were affiliated to *Flavobacterium columnare* and *Niastella koreensis*, which were also the most abundant *nosZ* taxa according to the metagenome analyses. Discrepancies between the amplicon sequencing data and the shotgun metagenome data were observed for NosZG2 and NosZG5 to some degree. The OTUs assigned to the genus *Thauera* and *Ruvrivivax gelatinosus* were the dominant among NosZG2, constituting 50.4% of the amplicons, but together constituted only 2.2% of the metagenome-derived *nosZ* sequences belonging to this group. Instead, an OTU affiliated to *Ramlibacter tataouinensis*, closely related to *R. gelatinosus*, was recovered in high relative abundance (15.4%) in the metagenome-derived *nosZ* pool. Likewise, the low relative abundance of the OTUs affiliated to *Dechloromonas aromatica* (0.2%) in the NosZG5 amplicons was coupled to the high relative abundance of the *nosZ* OTU affiliated to the *Azospira oryzae* (54.6%) with >96% translated amino acid identity within the amplified region. Thus, the observed discrepancy may be due to ambiguous OTU assignment among the closely related taxa.

The TaqMan-based qPCR assays were also compared with the most frequently used qPCR assays for clade I and clade II *nosZ* using SYBR green detection chemistry (Figure S6). Despite the low amplification efficiency (79.3%), the abundances of clade I *nosZ*, as determined with the SYBR Green assays, were relatively consistent with the cumulative abundances of NosZG1 and NosZG2 determined with the TaqMan qPCR assays, except for Daejeon1 sample, where the SYBR Green qPCR yielded 87 times lower copy numbers than the TaqMan qPCR. The SYBR Green qPCR with nosZII-F–nosZII-R primer set, with a subpar amplification efficiency of 61.9%, underestimated the clade II *nosZ* copy numbers by several orders of magnitudes for three of the four analyzed samples, with the largest difference (1.8 x 10^3^-fold) observed with the Daejeon1 sample. The importance of the clade II *nosZ* among the N_2_O-reducing microbial community in these activated sludge samples would have gone unnoticed due to this underestimation, if the conventional qPCR assays were used as the sole means of *nosZ* quantification.

### Quantitative analyses of *nosZ* genes and transcripts in activated sludge microbiomes

With the new group-specific *nosZ* primers, the activated sludge microbiomes from anoxic tanks of six A2O WWTPs apart from those used for development and cross-checking processes were analyzed for the *nosZ* gene and transcript abundances (Figure 4). As observed with the samples from Daejeon1, Daejeon2, Gwangju, and Gapyeong, the clade II *nosZ* (amplified with NosZG3 and NosZG5), were at least three-fold more abundant than the clade I *nosZ* (amplified with NosZG1 and NosZG2) in terms of gene abundance. The *Dechloromonas*-like *nosZ* genes (NosZG5) was the most abundant *nosZ* group in all samples but from Busan WWTP, where the abundance of *Flavobacterium*-like *nosZ* genes (NosZG3) was statistically similar to this group. The transcript profiles further highlighted the significance of clade II *nosZ* in activated sludge microbiomes. Due to the lower gene abundance and generally low level of transcription observed for clade I *nosZ* (transcript-to-gene ratios of <1.0 for both NosZG1 and NosZG2 in five out of six samples), transcription of the clade I *nosZ* was at least an order of magnitude lower than that of the clade II *nosZ* in all samples. The *Pseudomonas*-like *nosZ* (NosZG1) had remarkably low transcription level (transcript-to-gene ratios of <0.01) in all samples but from Busan WWTP, suggesting irrelevance of this *nosZ* group as an N_2_O sink in the activated sludge tanks. The relative importance of NosZG3 and NosZG5 is difficult to fathom, as *nosZ* transcripts targeted by NosZG3 were significantly more abundant in the sample from Busan WWTP (*p*<0.05), while NosZG5 was significantly more abundant in the samples from three other WWTPs (*p*<0.05).

**Figure 4.**
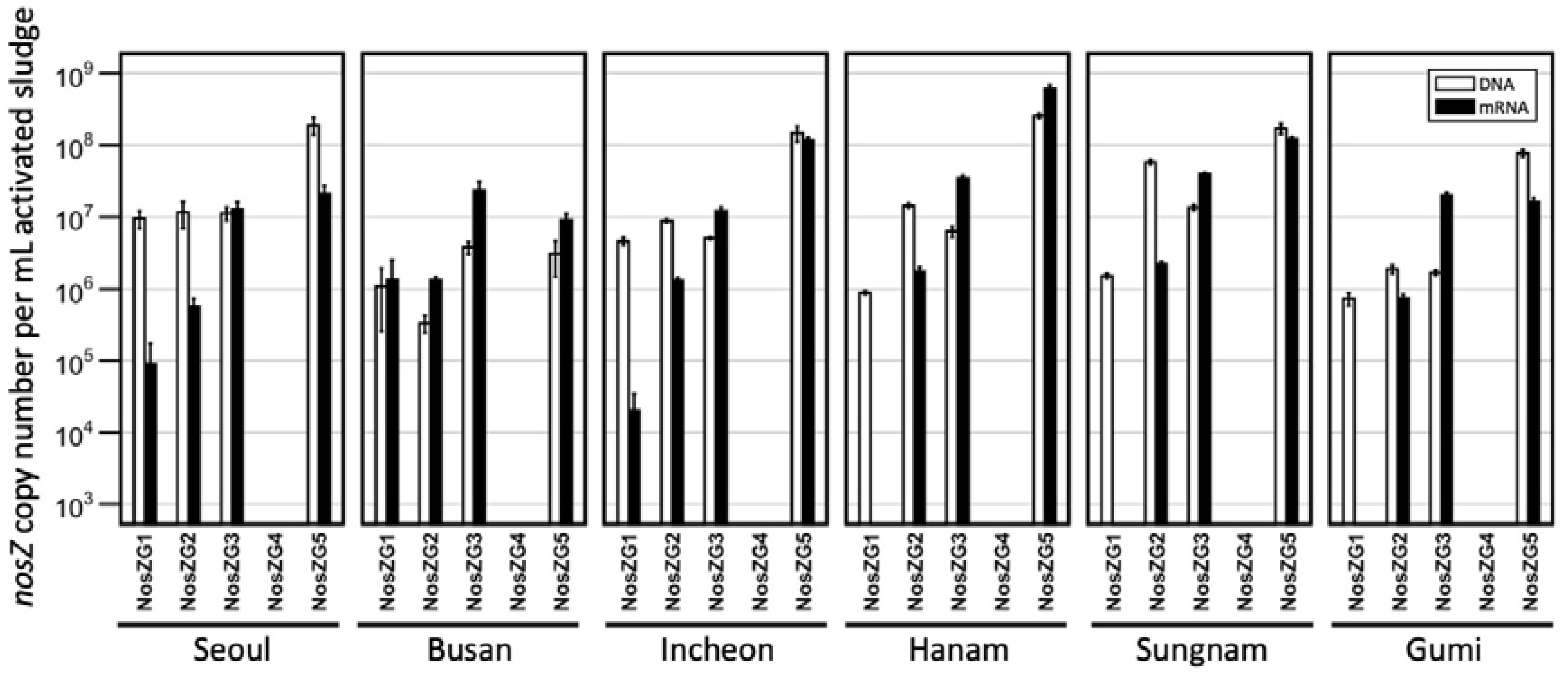
The copy numbers of NosZG1-5 *nosZ* genes and transcripts in activated sludge samples collected from anoxic tanks of six activated sludge-type WWTPs in Korea, as quantified using the group-specific *nosZ* qPCR developed in this study. The presented data are the averages of triplicate samples processed seperately through extraction, purification and reverse transcription (for analyses of transcripts), and qPCR procedures, with the error bars representing their standard deviations.

## Discussion

Quantification of the functional genes encoding the nitrogen cycle enzymes (e.g., bacterial/archaeal *amoA*, comammox *amoA*, *nrfA*, and clade I and II *nosZ*) has been used in increasing number of studies across diverse disciplines of environmental sciences and engineering, as predictors of nitrogen cycling activity in diverse soil and aquatic environments.^43–46^ Despite the frequent use of qPCR in determining the potential N_2_O-reducing populations, i.e., clade I and clade II *nosZ*-posessing microorganisms, the predictability of qPCR quantification has rarely been scrutinized in the previous studies. The new group-specific qPCR assays exhibited definite advantages over these frequently used primers. Although the targets were limited to the groups of *nosZ*-possessing organisms enriched with N_2_O in *ex situ* cultures, the copy numbers obtained were, in many cases, orders of magnitudes larger than those measured using previous primer sets. The diversity of the sequences recovered in the amplicon sequence indicates the exhaustive coverages of the qPCR assays developed in this study within the targeted groups. Most of the major *nosZ* OTUs (>1% in relative abundance) recovered in metagenome-based *nosZ* profiling of the examined activated sludge samples were amplified with exactly one of the four primer sets, thus confirming the absolute mutual exclusiveness of the primers/probe sets. Any methods for quantification of functional genes are prone to error, and the *nosZ* qPCR methods developed here also have certain deficiencies, including inability to amplify *nosZ* genes affiliated to relatively abundant *Iganvibacterium* spp. In designing the group-specific primers, a trade-off between coverage and mutual exclusivity was inevitable, which could have been the cause of modest discrepancy with the metagenome data. Nevertheless, despite these drawbacks, there is little doubt that the qPCR assays developed in this study are the most reliable tools for real-time quantification of the environmentally significant *nosZ* genes available to date.

The clade II *nosZ*-possessing organisms of the *Flavobacterium* (amplified with NosZG3), *Dechloromonas*, and *Azospira* (amplified with NosZG5) genera have previously been characterized with high-affinity reduction of N_2_O.^13, 14^ The whole-cell half-saturation constants as low as 0.324 μM has been reported for these groups of N_2_O reducers. The *nosZ* gene and transcript pools of all six of the activated sludge samples were dominated by these clade II *nosZ* (targeted by NosZG3 and NosZG5). This observation, along with the high transcript-to-gene ratios support the previous hypothesis that the clade II *nosZ*, with higher affinity to N_2_O, may have consequential role in reducing N_2_O emissions from soil and aquatic environments, including the activated sludge tanks.^15, 47^ The low transcript-to-gene ratios of the clade I *nosZ* (<0.1 in five out of six samples) targeted by NosZG1 and NosZG2 were also notable in the *nosZ* transcript profiles. The organisms possessing clade I *nosZ* genes, being almost exclusively denitrifiers, may favor upstream steps of denitrification reaction in the anoxic activated sludge tanks, where NO_3_^-^ and/or NO_2_^-^ are constantly available at millimolar concentrations and N_2_O is immediately consumed by their high-affinity cohabitants, which may prefer N_2_O to NO_3_^-^ or NO_2_^-^. More physiological evidences are warranted in the future research, however, to verify whether such hypothetical ‘division of labor’ really exists, as any further extrapolation from the *nosZ* transcription data alone would be an over-interpretation.

The major *nosZ*-possessing organisms identified in the qPCR and metagenomic analyses differ greatly from the major *nosZ*-possessing populations of the agricultural soils analyzed previously with shotgun metagenome sequencing.^48^ None of the six most abundant *nosZ* phylogenetic groups in the Havana and Urbana agricultural research site soils (the *nosZ* genes affiliated to *Anaeromyxobacter, Opitutus*, *Hydrogenobacter*, *Ignavibacterium*, *Dyedobacter*, and *Gemmatimonas*) was identified as a major population in any of the activated sludge samples examined in this study. Neither were they enriched with N_2_O to high relative abundance (e.g., >1%) in the fed-batch reactor. Of the organisms affiliated to these genera, *Anaeromyxobacter dehalogenans* and *Gemmatimonas aurentiaca* have been confirmed of N_2_O reduction activity; however, the N_2_O reduction rates measured *in vitro* were orders of magnitudes lower for these organisms than other examined N_2_O-reducing organisms throughout the entire range of N_2_O concentration, and *G. aurantiaca* lacked the capability to utilize N_2_O as the growth substrate.^11, 13, 30^ The differences in the compositions of the *nosZ*-possessing populations may be attributed to the inherent difference in the rates of the biochemical turnover processes in WWTP activated sludges and soils. Rapid nitrogen cycling reactions take place on a constant basis in WWTPs, while the time scale of the biogeochemical processes in agricultural soils is orders of magnitude longer and often, nitrogen supply provided through fertilization and plant exudation is sparse and sporadic.^49^ Besides, upland agricultural soils are often well-aerated and thus, the capability of N_2_O utilization would not provide specific benefits to the *nosZ*-harboring organisms at ordinary dry conditions.^50^ In fact, incubation of soil microbial consortium with N_2_O in the same fed-batch cultivation resulted in enrichment of the same organismal groups as those observed in the activated sludge enrichments (e.g., *Pseudomonas*, *Flavobacterium*, *Acidovorax,* and *Chryseobacterium*), suggesting that under occasions of pulse stimulation of denitrification and N_2_O reduction, e.g., flooding events immediately following fertilization, these organisms may become the relevant N_2_O sinks in soils, as well (Table S8). Besides, none of the 14 *nosZ*-carrying genera enriched with N_2_O in this study from either the activated sludge and soil belonged to the taxa identified as non-denitrifying N_2_O reducers suggesting that the abundance of non-denitrifier taxa, *per se*, may not be a measure of the N_2_O sink capability, as previously proposed.

## Acknowledgements

This work was financially supported by “the R&D Center for reduction of Non-CO2 Greenhouse Gases (Grant No. 2017002420002)” funded by Korea Ministry of Environment (MOE).

## Reference

1. Ciais, P.; Sabine, C.; Bala, G.; Bopp, L.; Brovkin, V.; Canadell, J.; Chhabra, A.; DeFries, R.; Galloway, J.; Heimann, M.; Jones, C.; Le Quere, C.; Myneni, R. B.; Piao, S.; Thornton, P., Carbon and other biogeochemical cycles. In Climate change 2013: The physical science basis. Contribution of working group I to the fifth assessment report of the Intergovernmental Panel on Climate Change, Stocker, T. F.; Qin, D.; Plattner, G.-K.; Tignor, M.; Allen, S. K.; Boschung, J.; Nauels, A.; Xia, Y.; Bex, V.; Midgley, P. M., Eds. Cambridge University Press: Cambridge, United Kingdom, New York, NY, USA, 2013; pp 465–570. http://www.ipcc.ch/pdf/assessment-report/ar5/wg1/WG1AR5_Chapter06_FINAL.pdf

2. Ravishankara, A. R.; Daniel, J. S.; Portmann, R. W. Nitrous oxide (N_2_O): the dominant ozone-depleting substance emitted in the 21^st^ century. Science 2009, 326, (5949), 123–125.

3. Portmann, R. W.; Daniel, J. S.; Ravishankara, A. R. Stratospheric ozone depletion due to nitrous oxide: influences of other gases. Phil. Trans. R. Soc. B 2012, 367, (1593), 1256–1264.

4. Yoon, S.; Song, B.; Phillips, R. L.; Chang, J.; Song, M. J. Ecological and physiological implications of nitrogen oxide reduction pathways on greenhouse gas emissions in agroecosystems. FEMS Microbiol. Ecol. 2019, 95, (6), fiz066.

5. Law, Y.; Ye, L.; Pan, Y.; Yuan, Z. Nitrous oxide emissions from wastewater treatment processes. Phil. Trans. R. Soc. B 2012, 367, (1593), 1265–1277.

6. Thomson, A. J.; Giannopoulos, G.; Pretty, J.; Baggs, E. M.; Richardson, D. J. Philos Trans Royal Soc B 2012, 367, (1593), 1157–1168.

7. Frutos, O. D.; Quijano, G.; Aizpuru, A.; Muñoz, R. A state-of-the-art review on nitrous oxide control from waste treatment and industrial sources. Biotechnol. Adv. 2018, 36, (4), 1025–1037.

8. Hallin, S.; Philippot, L.; Löffler, F. E.; Sanford, R. A.; Jones, C. M. Genomics and ecology of novel N_2_O-reducing microorganisms. Trends Microbiol. 2017, 26, (1), 43–55.

9. Suenaga, T.; Hori, T.; Riya, S.; Hosomi, M.; Smets, B. F.; Terada, A. Enrichment, isolation, and characterization of high-affinity N_2_O-reducing bacteria in a gas-permeable membrane reactor. Environ. Sci. Technol. 2019, in press; DOI: 10.1021/acs.est.9b02237

10. Zumft, W. G. Cell biology and molecular basis of denitrification. Microbiol. Mol. Biol. Rev. 1997, 61, (4), 533–616.

11. Sanford, R. A.; Wagner, D. D.; Wu, Q.; Chee-Sanford, J. C.; Thomas, S. H.; Cruz-García, C.; Rodríguez, G.; Massol-Deyá; Krishnani, K. K.; Ritalahti, K. M.; Nissen, S.; Konstantinidis, K. T.; Löffler, F. E. Unexpected nondenitrifier nitrous oxide reductase gene diversity and abundance in soils. Proc. Natl Acad Sci 2012, 109, (48), 19709–19714.

12. Conthe, M.; Wittorf, L.; Kuenen, J. G.; Kleerebezem, R.; van Loosdrecht, M. C. M.; Hallin, S. Life on N_2_O: deciphering the ecophysiology of N_2_O respiring bacterial communities in a continuous culture. ISME J. 2018, 12, (4), 1142–1153.

13. Yoon, S.; Nissen, S.; Park, D.; Sanford, R. A.; Löffler, F. E. Nitrous oxide reduction kinetics distinguish bacteria harboring clade I versus clade II NosZ. Appl. Environ. Microbiol. 2016, 82, (13),3793–3800.

14. Suenaga, T.; Riya, S.; Hosomi, M.; Terada, A. Biokinetic characterization and activities of N_2_O-reducing bacteria in response to various oxygen levels. Front. Microbiol. 2018, 9, 697.

15. Jones, C. M.; Spor, A.; Brennan, F. P.; Breuil, M.-C.; Bru, D.; Lemanceau, P.; Griffiths, B.; Hallin, S.; Philippot, L. Recently identified microbial guild mediates soil N_2_O sink capacity. *Nat*. Clim. Change 2014, 4, 801–805.

16. Domeignoz-Horta, L. A.; Spor, A.; Bru, D.; Breuil, M.-C.; Bizouard, F.; Leonard, J.; Philippot, L. The diversity of the N_2_O reducers matters for the N_2_O:N_2_ denitrification end-product ratio across an annual and a perennial cropping system. Front. Microbiol. 2015, 6, 971

17. Yoon, H.; Song, M. J.; Yoon, S. Design and feasibility analysis of a self-sustaining biofiltration system for removal of low concentration N_2_O emitted from wastewater treatment plants. Environ. Sci. Technol. 2017, 51, (18), 10736–10745.

18. Yoon, H.; Song, M. J.; Kim, D. D.; Sabba, F.; Yoon, S. A serial biofiltration system for effective removal of low-concentration nitrous oxide in oxic gas streams: mathematical modeling of reactor performance and experimental validation. Environ. Sci. Technol. 2019, 53, (4), 2063–2074.

19. Paranychianakis, N. V.; Tsiknia, M.; Kalogerakis, N. Pathways regulating the removal of nitrogen in planted and unplanted subsurface flow constructed wetlands. Water Res. 2016, 102, 321–329.

20. Conthe, M.; Kuenen, J. G.; Kleerebezem, R.; van Loosdrecht, M. C. M. Exploring microbial N_2_O reduction: a continuous enrichment in nitrogen free medium. Environ. Microbiol. Rep. 2018, 10, (1), 102–107.

21. Boonnorat, J.; Techkarnjanaruk, S.; Honda, R.; Ghimire, A.; Angthong, S.; Rojviroon, T.; Phanwilai, S. Enhanced micropollutant biodegradation and assessment of nitrous oxide concentration reduction in wastewater treated by acclimatized sludge bioaugmentation. Sci. Tot. Environ. 2018, 637, 771–779.

22. Song, K.; Suenaga, T.; Harper, W. F.; Hori, T.; Riya, S.; Hosomi, M.; Terada, A. Effects of aeration and internal recycle flow on nitrous oxide emissions from a modified Ludzak–Ettinger process fed with glycerol. Environ. Sci. Pollut. Res. 2015, 22, (24), 19562–19570.

23. Vieira, A.; Galinha, C. F.; Oehmen, A.; Carvalho, G. The link between nitrous oxide emissions, microbial community profile and function from three full-scale WWTPs. Sci. Tot. Environ. 2019, 651, 2460–2472.

24. Henry, S.; Bru, D.; Stres, B.; Hallet, S.; Philippot, L. Quantitative detection of the *nosZ* gene, encoding nitrous oxide reductase, and comparison of the abundances of 16S rRNA, *narG*, *nirK*, and *nosZ* genes in soils. Appl. Environ. Microbiol. 2006, 72, (8), 5181–5189.

25. Jones, C. M.; Graf, D. R.; Bru, D.; Philippot, L.; Hallin, S. The unaccounted yet abundant nitrous oxide-reducing microbial community: a potential nitrous oxide sink. ISME J. 2013, 7, (2), 417–426.

26. Di, H. J.; Cameron, K. C.; Podolyan, A.; Robinson, A. Effect of soil moisture status and a nitrification inhibitor, dicyandiamide, on ammonia oxidizer and denitrifier growth and nitrous oxide emissions in a grassland soil. Soil Biol. Biochem. 2014, 73, 59–68.

27. Stoliker, D. L.; Repert, D. A.; Smith, R. L.; Song, B.; LeBlanc, D. R.; McCobb, T. D.; Conaway, C. H.; Hyun, S. P.; Koh, D.-C.; Moon, H. S. Hydrologic controls on nitrogen cycling processes and functional gene abundance in sediments of a groundwater flow-through lake. Environ. Sci. Technol. 2016, 50, (7), 3649–3657.

28. Binder, B. J.; Liu, Y. C., Growth rate regulation of rRNA content of a marine *Synechococcus* (Cyanobacterium) strain. Appl. Environ. Microbiol. 1998, 64, (9), 3346–3351.

29. Yoon, S.; Sanford, R. A.; Löffler, F. E., *Shewanella* spp. use acetate as an electron donor for denitrification but not ferric iron or fumarate reduction. Appl. Environ. Microbiol. 2013, 79, (8), 2818–2822.

30. Park, D.; Kim, H.; Yoon, S. Nitrous oxide reduction by an obligate aerobic bacterium *Gemmatimonas aurantiaca* strain T-27. Appl. Environ. Microbiol. 2017, 83, (12), e00502–17.

31. Ritalahti, K. M.; Amos, B. K.; Sung, Y.; Wu, Q.; Koenigsberg, S. S.; Löffler, F. E. Quantitative PCR targeting 16S rRNA and reductive dehalogenase genes simultaneously monitors multiple *Dehalococcoides* strains. Appl. Environ. Microbiol. 2006, 72, (4), 2765–2774.

32. Edgar, R. C. MUSCLE: multiple sequence alignment with high accuracy and high throughput. Nucleic Acids Res. 2004, 32, (5), 1792–1797.

33. Kumar, S.; Stecher, G.; Tamura, K. MEGA7: Molecular Evolutionary Genetics Analysis version 7.0 for bigger datasets. Mol. Biol. Evol. 2016, 33, (7), 1870–1874.

34. Hysom, D. A.; Naraghi-Arani, P.; Elsheikh, M.; Carrillo, A. C.; Williams, P. L.; Gardner, S. N. Skip the alignment: degenerate, multiplex primer and probe design using K-mer matching instead of alignments. PloS One 2012, 7, (4), e34560.

35. Gardner, S. N.; Slezak, T. J. B. B. Simulate_PCR for amplicon prediction and annotation from multiplex, degenerate primers and probes. BMC Bioinformatics 2014, 15, (1), 237.

36. Yoon, S.; Cruz-Garcia, C.; Sanford, R.; Ritalahti, K. M.; Loffler, F. E. Denitrification versus respiratory ammonification: environmental controls of two competing dissimilatory NO_3_^−^/NO_2_^−^ reduction pathways in *Shewanella loihica* strain PV-4. ISME J. 2015, 9, (5), 1093–1104.

37. Li, W.; Godzik, A. Cd-hit: a fast program for clustering and comparing large sets of protein or nucleotide sequences. Bioinformatics 2006, 22, (13), 1658–1659.

38. Bolger, A. M.; Lohse, M.; Usadel, B. Trimmomatic: a flexible trimmer for Illumina sequence data. Bioinformatics 2014, 30, (15), 2114–2120.

39. Nurk, S.; Meleshko, D.; Korobeynikov, A.; Pevzner, P. A. metaSPAdes: a new versatile metagenomic assembler. Genome Res. 2017, 27, (5), 824–834.

40. Li, H.; Handsaker, B.; Wysoker, A.; Fennell, T.; Ruan, J.; Homer, N.; Marth, G.; Abecasis, G.; Durbin, R. The sequence alignment/map format and SAMtools. Bioinformatics 2009, 25, (16), 2078–2079.

41. Miller, C. S.; Baker, B. J.; Thomas, B. C.; Singer, S. W.; Banfield, J. F. EMIRGE: reconstruction of full-length ribosomal genes from microbial community short read sequencing data. Genome Biol. 2011, 12, (5), R44.

42. Huang, Y.; Gilna, P.; Li, W. Identification of ribosomal RNA genes in metagenomic fragments. Bioinformatics 2009, 25, (10), 1338–1340.

43. Pjevac, P.; Schauberger, C.; Poghosyan, L.; Herbold, C. W.; van Kessel, M. A. H. J.; Daebeler, A.; Steinberger, M.; Jetten, M. S. M.; Lücker, S.; Wagner, M.; Daims, H. *amoA*-targeted polymerase chain reaction primers for the specific detection and quantification of comammox *Nitrospira* in the environment. Front. Microbiol. 2017, 8, 1508.

44. Meinhardt, K. A.; Bertagnolli, A.; Pannu, M. W.; Strand, S. E.; Brown, S. L.; Stahl, D. Evaluation of revised polymerase chain reaction primers for more inclusive quantification of ammonia-oxidizing archaea and bacteria. Environ. Microbiol. Rep. 2015, 7, (2), 354–363.

45. Ma, Y.; Zilles, J. L.; Kent, A. D. An evaluation of primers for detecting denitrifiers via their functional genes. Environ Microbiol 2019, 21, (4), 1196–1210.

46. Chen, X.; Peltier, E.; Sturm, B. S. M.; Young, C. B. Nitrogen removal and nitrifying and denitrifying bacteria quantification in a stormwater bioretention system. Water Res. 2013, 47, (4), 1691–1700.

47. Conthe, M.; Wittorf, L.; Kuenen, J. G.; Kleerebezem, R.; Hallin, S.; van Loosdrecht, M. C. M. Growth yield and selection of *nosZ* clade II types in a continuous enrichment culture of N_2_O respiring bacteria. Environ. Microbiol. Rep. 2018, 10, (3), 239–244.

48. Orellana, L. H.; Rodriguez-R, L. M.; Higgins, S.; Chee-Sanford, J. C.; Sanford, R. A.; Ritalahti, K. M.; Löffler, F. E.; Konstantinidis, K. T. Detecting nitrous oxide reductase (*nosZ*) genes in soil metagenomes: method development and implications for the nitrogen cycle. mBio 2014, 5, (3), e01193–14.

49. Cassman, K. G.; Dobermann, A.; Walters, D. T. Agroecosystems, nitrogen-use efficiency, and nitrogen management. Ambio 2002, 31, (2), 132–140.

50. Aulakh, M. S.; Khera, T. S.; Doran, J. W. Mineralization and denitrification in upland, nearly saturated and flooded subtropical soilI. Effect of nitrate and ammoniacal nitrogen. Biol. Fert. Soils 2000, 31, (2), 162–167.

