## Supplementary material for "Quantification of *nosZ* genes and transcripts in activated sludge microbiomes with novel group-specific qPCR methods validated with metagenomic analyses": Supplmental material

**Running Head:** Quantification of *nosZ* with group-specific qPCR

*Present address: Department of Civil and Environmental Engineering, Georgia Institute of Technology, Atlanta, Georgia, USA

†Present address: Department of Civil and Environmental Engineering, Cornell University, Ithaca, New York, USA

**Summary of Content:**

**Total Pages: 20**

**8 Tables, 6 Figures**

**Microbial community analysis of the batch culture fed with N_2_O as a sole electron acceptor**

The 16S rRNA amplicon sequences generated through Illumina Miseq platform were processed using QIIME pipeline v 1.9.1. The sequence reads with the quality score lower than the default cut-off (phred score 20) were removed. The trimmed reads were clustered to the Greengenes v 13.8 16S rRNA gene database with the cut-off value set to 0.97. The remaining reads, which failed to cluster with the reference database, were clustered into *de novo* OTUs with the same cuff-off value (0.97). The taxonomy was assigned to each OTU using the RDP classifier against the Greengenes v 13.8 database.

Table S1. The 174 *nosZ* sequences of the 14 genera identified as substantially enriched (to >0.3% relative abundance) organisms after fed-batch incubation on N_2_O.

| Group | Organism | Amino acid sequence  accession number | Nucleotide sequence  accession number | Assembly  level |
| --- | --- | --- | --- | --- |
| NosZG1 | *Rhodobacter capsulatus* SB 1003 | ADE87331 | CP001313 | Complete |
|  | *Rhodobacter* sp. AKP1 | EKX57490 | ANFS01000017 | Draft |
|  | *Rhodobacter* sp. CACIA14H1 | ESW60705 | AYNO01000067 | MAG |
|  | *Rhodobacter johrii* | ODM44753 | MABH01000036 | Draft |
|  | *Pseudomonas stutzeri* | CAA37714 | X53676 | Clone |
|  | *Pseudomonas aeruginosa* | CAA46381 | X65277 | Clone |
|  | *Pseudomonas aeruginosa* PAO1 | AAG06780 | AE004091 | Complete |
|  | *Pseudomonas fluorescens* | AAG34386 | AF197468 | Clone |
|  | *Pseudomonas stutzeri* | AEJ06803 | CP002881 | Complete |
|  | *Pseudomonas stutzeri* TS44 | EIK51269 | AJXE01000042 | Draft |
|  | *Pseudomonas stutzeri* 19SMN4 | AHY41575 | CP007509 | Complete |
|  | *Pseudomonas stutzeri* A1501 | ABP81178 | CP000304 | Complete |
|  | *Pseudomonas stutzeri* ODKF13 | KXO84969 | LSVE01000001 | Draft |
|  | *Pseudomonas stutzeri* DSM 4166 | AEA85507 | CP002622 | Complete |
|  | *Pseudomonas stutzeri* ST-9 | KKJ94548 | JXJL01000037 | Draft |
|  | *Pseudomonas stutzeri* | KJH80175 | JYHV01000034 | Draft |
|  | *Pseudomonas stutzeri* | AKN28567 | CP011854 | Complete |
|  | *Pseudomonas stutzeri* YC-YH1 | KIL03776 | JUDR01000006 | Draft |
|  | *Pseudomonas stutzeri* 273 | ANF25877 | CP015641 | Complete |
|  | *Pseudomonas stutzeri* CCUG 29243 | AFM31993 | CP003677 | Complete |
|  | *Pseudomonas stutzeri* RCH2 | AGA85386 | CP003071 | Complete |
|  | *Pseudomonas stutzeri* B1SMN1 | EPL64545 | AMVM01000001 | Draft |
|  | *Pseudomonas stutzeri* TS44 | EIK53500 | AJXE01000006 | Draft |
|  | *Pseudomonas stutzeri* NF13 | EME01589 | AOBS01000021 | Draft |
|  | *Pseudomonas stutzeri* BAL361 | KIZ35892 | JXXD01000101 | Draft |
|  | *Pseudomonas stutzeri* KOS6 | EWC39540 | AMCZ02000036 | Draft |
|  | *Pseudomonas stutzeri* | KZX57309 | LWEF01000296 | Draft |
|  | *Pseudomonas flexibilis* | KHL68146 | JRUD01000026 | Draft |
|  | *Pseudomonas bauzanensis* | EZQ19619 | JFHS01000001 | Draft |
|  | *Pseudomonas* sp. BAY1663 | EXF45396 | AZSV01000025 | Draft |
|  | *Pseudomonas stutzeri* MF28 | EQM73584 | ATAR01000051 | Draft |
|  | *Pseudomonas aeruginosa* UCBPP-PA14 | ABJ12646 | CP000438 | Complete |
|  | *Pseudomonas aeruginosa* str. Stone 130 | EMZ56463 | AQFN01000013 | Draft |
|  | *Pseudomonas aeruginosa* PA7 | ABR85628 | CP000744 | Complete |
|  | *Pseudomonas saudimassiliensis* | CEA03480 | LM997413 | Draft |
|  | *Pseudomonas mandelii* PD30 | KDD69024 | AZQQ01000074 | Draft |
|  | *Pseudomonas citronellolis* | AMO73766 | CP014158 | Complete |
|  | *Pseudomonas aeruginosa* DK1 | CRZ29134 | LN870292 | Complete |
|  | *Pseudomonas xanthomarina* | CEG55166 | CCYE01000056 | Draft |
|  | *Pseudomonas fluorescens* C3 | KJZ46623 | LACD01000005 | Draft |
|  | *Pseudomonas* sp. HMP271 | KGK83321 | JMFZ01000007 | Draft |
|  | *Pseudomonas* sp. BICA1-14 | KJS72355 | JUEC02000100 | MAG |
|  | *Pseudomonas brassicacearum* subsp. brassicacearum NFM421 | AEA68359 | CP002585 | Complete |
|  | *Pseudomonas brassicacearum* subsp. brassicacearum NFM421 | AEA68936 | CP002585 | Complete |
|  | *Pseudomonas* sp. AAC | KES23648 | JNCW01000008 | Draft |
|  | *Pseudomonas* sp. Chol1 | EKM97315 | AMSL01000021 | Draft |
|  | *Pseudomonas stutzeri* TR2 | BAM68548 | AB764137 | Clone |
|  | *Pseudomonas mendocina* NK-01 | AEB56170 | CP002620 | Complete |
|  | *Pseudomonas* sp. P179 | EMZ56821 | AQFO01000015 | Draft |
|  | *Pseudomonas* sp. HMP271 | KGK82459 | JMFZ01000010 | Draft |
|  | *Pseudomonas fluorescens* F113 | AEV63001 | CP003150 | Complete |
|  | *Pseudomonas* sp. K35 | OCX95855 | MCAL01000135 | MAG |
|  | *Pseudomonas* sp. ATCC 13867 | AGI26824 | CP004143 | Complete |
|  | *Pseudomonas mandelii* JR-1 | AHZ73475 | CP005961 | Complete |
|  | *Pseudomonas fluorescens* Q2-87 | EJL01169 | AGBM01000001 | Draft |
|  | *Pseudomonas extremaustralis 14-3* substr. 14-3b | EZI26062 | AHIP01000029 | Draft |
|  | *Pseudomonas* sp. TTU2014-105ASC | KRW65887 | LKKM01000027 | Draft |
|  | *Pseudomonas saudiphocaensis* | CDZ95800 | CCSF01000001 | Draft |
|  | *Pseudomonas fluorescens* NCIMB 11764 | AKV08877 | CP010945 | Complete |
|  | *Pseudomonas syringae* CEB003 | KFE56324 | JPQT01000010 | Draft |
|  | *Pseudomonas mandelii* PD30 | KDD65257 | AZQQ01000110 | Draft |
|  | *Pseudomonas syringae* Riq4 | KNH28443 | LFQK01000013 | Draft |
|  | *Pseudomonas veronii* 1YdBTEX2 | SBW80421 | LT599583 | Complete |
|  | *Pseudomonas fluorescens* UM270 | KIQ58959 | JXNZ01000105 | Draft |
|  | *Pseudomonas veronii* DSM 11331 | KRP62004 | JYLL01000052 | Draft |
|  | *Pseudomonas* sp. FH4 | ETK17447 | AOHN01000020 | Draft |
|  | *Pseudomonas fluorescens* NT0133 | KJH85519 | JYHW01000047 | Draft |
|  | *Pseudomonas fluorescens* C1 | KJZ39917 | LACE01000004 | Draft |
|  | *Pseudomonas fluorescens* C3 | KJZ44076 | LACD01000014 | Draft |
|  | *Pseudomonas fluorescens* H24 | KJZ65387 | LACH01000022 | Draft |
|  | *Pseudomonas xanthomarina strain* UASWS0955 | OCX24688 | MDEM01000020 | Draft |
|  | *Pseudomonas fluorescens* C1 | KJZ39900 | LACE01000004 | Draft |
|  | *Pseudomonas fluorescens* FW300-N2C3 | ALI10524 | CP012831 | Complete |
|  | *Pseudomonas brenneri* RGCB | OAE15347 | LVWZ01000025 | Draft |
|  | *Pseudomonas corrugata* RM1-1-4 | AOE60894 | CP014262 | Complete |
|  | *Pseudomonas fluorescens* C1 | KJZ39573 | LACE01000005 | Draft |
|  | *Pseudomonas chlororaphis* UFB2 | AKJ99839 | CP011020 | Complete |
|  | *Pseudomonas fluorescens* F113 | AEV63425 | CP003150 | Complete |
|  | *Pseudomonas fluorescens* FW300-N2E2 | AMZ70880 | CP015225 | Complete |
|  | *Pseudomonas lini* DSM 16768 | KMM89163 | JYLB01000010 | Draft |
|  | *Pseudomonas fluorescens* FW300-N1B4 | KZN21032 | LUKJ01000001 | Draft |
|  | *Pseudomonas brassicacearum* DF41 | AHL34282 | CP007410 | Complete |
|  | *Pseudomonas brassicacearum* DF41 | AHL34136 | CP007410 | Complete |
|  | *Pseudomonas* sp. TKP | AHC35295 | CP006852 | Complete |
|  | *Pseudomonas brassicacearum* DF41 | AHL34765 | CP007410 | Complete |
|  | *Pseudomonas bauzanensis* W13Z2 | EZQ12858 | JFHS01000040 | Draft |
|  | *Pseudomonas* sp. GM41 | EUB74376 | AKJN02000006 | Draft |
|  | *Pseudomonas* sp. KG01 | KMT52756 | LFMW01000022 | Draft |
|  | *Pseudomonas* sp. AU11447 | OBY93498 | LZDG02000013 | Draft |
|  | *Pseudomonas* sp. AU12215 | OBY60969 | MACM02000005 | Draft |
|  | *Pseudomonas* sp. GM60 | EJM79615 | AKJI01000086 | Draft |
|  | *Pseudomonas* sp. GM67 | EJM91426 | AKJH01000023 | Draft |
|  | *Pseudomonas kilonensis* 1855-344 | KKA09578 | JZXC01000002 | Draft |
|  | *Pseudomonas thivervalensis* LMG 21626 | OAB49649 | LRSO01000023 | Draft |
|  | *Pseudomonas* sp. LZ-4 | ANT47466 | KU192988 | Clone |
|  | *Pseudomonas* sp. TTU2014-066ASC | KRW67808 | LKKJ01000021 | Draft |
|  | *Pseudomonas* sp. CFII68 | EPJ95261 | ATLN01000042 | Draft |
|  | *Pseudomonas* sp. 21 | KJJ98743 | JYOA01000007 | Draft |
|  | *Pseudomonas* sp. Root401 | KQW41490 | LMDO01000001 | Draft |
|  | *Pseudomonas* sp. CFII64 | EPJ87567 | ATLO01000011 | Draft |
|  | *Pseudomonas* sp. Root329 | KQV22503 | LMCV01000002 | Draft |
|  | *Pseudomonas silesiensis* A3 | ANJ56636 | CP014870 | Complete |
|  | *Pseudomonas* sp. Root401 | KQW30300 | LMDO01000018 | Draft |
|  | *Pseudomonas* sp. URMO17WK12:I11 | CRL52212 | LN854573 | Complete |
|  | *Pseudomonas litoralis* 2SM5 | SDS77717 | LT629748 | Complete |
|  | *Pseudomonas xanthomarina* LMG 23572 | SEH67047 | LT629970 | Complete |
|  | *Pseudomonas corrugata* BS3649 | SDU91569 | LT629798 | Complete |
|  | *Pseudomonas mucidolens* LMG 2223 | SDU92636 | LT629802 | Complete |
|  | *Pseudomonas pohangensis* DSM 17875 | SDU06131 | LT629785 | Complete |
|  | *Acinetobacter baumannii* | SCY30937 | FMVV01000007 | Draft |
| NosZG2 | *Acidovorax* sp. JS42 | ABM41372 | CP000539 | Complete |
|  | *Acidovorax* sp. MR-S7 | GAD21553 | DF238908 | Draft |
|  | *Acidovorax* sp. GW101-3H11 | KZT13955 | LUKZ01000028 | Draft |
|  | *Acidovorax* sp. SCN 68-22 | ODS70783 | MEDL01000013 | MAG |
|  | *Acidovorax* sp. RAC01 | AOG25129 | CP016447 | Complete |
|  | *Acidovorax delafieldii* 2AN | EER59730 | ACQT01000101 | Draft |
|  | *Acidovorax ebreus* TPSY | ACM32537 | CP001392 | Complete |
|  | *Acidovorax* sp. NO-1 | EHL20417 | AGTS01000159 | Draft |
|  | *Acidovorax* sp. SD340 | KQB61023 | LHUP01000064 | Draft |
|  | *Acidovorax* sp. Root70 | KRB42134 | LMHQ01000001 | Draft |
|  | *Acidovorax* sp. Root275 | KRD41873 | LMJH01000010 | Draft |
|  | *Acidovorax temperans* | KJA11266 | JXYQ01000018 | Draft |
|  | *Acidovorax* sp. SCN 65-108 | ODS66146 | MEDK01000019 | MAG |
|  | *Diaphorobacter* sp. J5-51 | KLR57518 | JSYI01000085 | Draft |
|  | *Massilia* sp. NR 4-1 | AKU20495 | CP012201 | Complete |
|  | *Massilia* sp. Root351 | KQV90309 | LMDJ01000002 | Draft |
|  | *Massilia* sp. Root418 | KQW93573 | LMEC01000020 | Draft |
|  | *Massilia violaceinigra* B2 | ATQ76468 | CP024608 | Complete |
|  | *Thauera* sp. SWB20 | KIN90102 | JTDM01000019 | Draft |
|  | *Thauera* sp. SWB20 | KIN91605 | JTDM01000013 | Draft |
|  | *Thauera linaloolentis* 47Lol = DSM 12138 | ENO89111 | AMXE01000020 | Draft |
|  | *Thauera* sp. 27 | ENO81470 | AMXB01000011 | Draft |
|  | *Thauera* sp. 63 | ENO75848 | AMXC01000025 | Draft |
|  | *Thauera humireducens* | AMO37672 | CP014646 | Complete |
|  | *Thauera terpenica* 58Eu | EPZ15520 | ATJV01000055 | Draft |
|  | *Thauera chlorobenzoica* | SEG11938 | FNVJ01000018 | Draft |
|  | *Thiobacillus denitrificans* ATCC 25259 | AAZ97342 | CP000116 | Complete |
|  | *Thiobacillus denitrificans* | KVW96019 | LDUG01000021 | Draft |
|  | *Thiobacillus* sp. SCN 64-317 | ODV13413 | MEGO01000014 | MAG |
|  | *Thiobacillus* sp. SCN 62-729 | ODU31360 | MEFL01000016 | MAG |
|  | *Thiobacillus* sp. SCN 63-57 | ODV04667 | MEGP01000002 | MAG |
|  | *Thiobacillus* sp. SCN 63-1177 | ODU00913 | MEFI01000158 | MAG |
| NosZG3 | *Chryseobacterium koreense* CCUG 49689 | KMQ71079 | LFNG01000011 | Draft |
|  | *Chryseobacterium antarcticum_*LMG_24720 | KEY18552 | JPEP01000002 | Draft |
|  | *Chryseobacterium jeonii_*DSM_17048 | KIA89615 | JSYL01000002 | Draft |
|  | *Chryseobacterium solincola* DSM_22468 | KIA84180 | JSYK01000002 | Draft |
|  | *Chryseobacterium* sp. SCN 40-13 | ODS87522 | MEDR01000038 | MAG |
|  | *Flavobacterium anhuiense* GSE09 | AOC96669 | CP016907 | Complete |
|  | *Flavobacterium aquatile* LMG 4008 = ATCC 11947 | KGD69011 | JRHH01000002 | Draft |
|  | *Flavobacterium* sp. Root186 | KRB55478 | LMHW01000013 | Draft |
|  | *Flavobacterium_piscis_* CCUG_60099_FLP11 | OCB76033 | LVEN01000011 | Draft |
|  | *Flavobacterium columnare* ATCC 49512 | AEW86926 | CP003222 | Complete |
|  | *Flavobacterium* sp. F52 | EJF99141 | AKZQ01000042 | Draft |
|  | *Flavobacterium cauense* R2A-7 | ESU19613 | AVBI01000016 | Draft |
|  | *Flavobacterium limnosediminis* JC2902 | ESU29215 | AVGG01000002 | Draft |
|  | *Flavobacterium saliperosum* S13 | ESU27709 | AVFO01000002 | Draft |
|  | *Flavobacterium suncheonense* GH29-5 = DSM 17707 | KGO89605 | JRLW01000008 | Draft |
|  | *Flavobacterium enshiense* DK69 | KGO96403 | JRLZ01000004 | Draft |
|  | *Flavobacterium* sp. 316 | KIX20325 | JYGZ01000006 | Draft |
|  | *Flavobacterium columnare* 94-081 | AMA49664 | CP013992 | Complete |
|  | *Flavobacterium_urumqiense _*CGMCC_1.9230 | SEG02668 | FNVP01000005 | Draft |
|  | *Cloacibacterium_normanense* NRS-1 | OEL12465 | MKGI01000004 | Draft |
| NosZG4 | *Ignavibacterium_album_*JCM_16511 | AFH48560 | CP003418 | Complete |
|  | *Ignavibacteria bacterium* CG2_30_36_16 | OIP62760 | MNYQ01000044 | MAG |
|  | *Ignavibacteria bacterium* GWC2_38_9 | OGU69616 | MHAA01000010 | MAG |
|  | *Ignavibacteria bacterium* GWA2_36_19 | OGU38146 | MGZR01000108 | MAG |
|  | *Ignavibacteria bacterium* CG1_02_37_35 | OIO21410 | MNVA01000053 | MAG |
|  | *Ignavibacteria bacterium* GWB2_35_6b | OGU35993 | MGZW01000068 | MAG |
|  | *Ignavibacteria bacterium* GWA2_35_8 | OGU14412 | MGZP01000075 | MAG |
| NosZG5 | *Dechloromonas denitrificans* ED1 | ALB05717 | KT592356 | Draft |
|  | *Dechloromonas_aromatica_*RCB | AAZ46320 | CP000089 | Complete |
|  | *Dechlorosoma_suillum_*PS *[Azospira oryzae]* | AEV25288 | CP003153 | Complete |
|  | *Thauera_phenylacetica_*B4P_Cont77 | ENO96869 | AMXF01000077 | Draft |
|  | *Thauera_linaloolentis_*47Lol_=_DSM_12138 | ENO89642 | AMXE01000011 | Draft |

Table S2. The primer and probe sequences and the reaction conditions for quantitative PCR performed in this study (in addition to the primer and probe sets shown in Table 1)

| **Target genes** | **Primer/probe** | **Sequence** | **Slope** | ***y*-intercept** | **Efficiency (%)** | **Reference** | **Thermal Cycling** |
| --- | --- | --- | --- | --- | --- | --- | --- |
| **Eubacterial 16S rRNA gene** | Bac1055YF | ATG GYT GTC GTC AGC T | -3.58 | 39.09 | 90.3 | Ritalahti *et al*., 2006 | (95°C, 10 min) x 1  (95°C, 30 s; 56 °C, 1 min; 72°C, 1min) x 40 |
|  | Bac1392R | ACG GGC GGT GTG TAC |  |  |  |  |  |
|  | Bac1115Probe | FAM-CAA CGA GCG CAA CCC-TAMRA |  |  |  |  |  |
| **nosZ I** | 1840F | CGC RAC GGC AAS AAG GTS MSS GT | -3.94 | 34.52 | 79.3 | Henry *et al*., 2006 | (95°C, 15 min) x 1  (95°C, 15 s;(65°C – 60°C, -1°/cycle), 30 s; 72°C, 30 s; 80°C, 15s) x6  (95°C, 15 s; 60°C, 15 s; 72°C, 30 s; 80°C, 15 s) x 40  (95°C, 15 s;(60 to 95° C, 10 s, increment 0.5°)), x 1 |
|  | 2090R | CAK RTG CAK SGC RTG GCA GAA |  |  |  |  |  |
| **nosZ II** | nosZII-F | CTI GGI CCI YTK CAY AC | -4.78 | 50.48 | 61.9 | Jones *et al*., 2013 | (95°C, 7 min) x 1  (95°C, 15 s; 54°C, 30 s; 72°C, 30 s; 80°C, 30 s) x 50  (95°C, 15 s;(60 to 95° C, 10 s, increment 0.5°)), x 1 |
|  | nosZII-R | GCI GAR CAR AAI TCB GTR C |  |  |  |  |  |
| **luciferase**  **control cDNA** | luc_refA | TAC AAC ACC CCA ACA TCT TCG A | -3.48 | 36.35 | 93.8 | Johnson et al., 2005 | (95°C, 10 min) x 1  (95°C, 15 s; 60°C, 1 min) x 40  (95°C, 15 s;(60 to 95° C, continuous increment)), x 1 |
|  | luc_refB | GGA AGT TCA CCG GCG TCA T |  |  |  |  |  |

Table S3. The list of *nosZ* sequences imported from the UniProt database for construction of the Hidden Markov Model (HMM) for *nosZ* clade I

| Accession No. | Organism | Length (aa) |
| --- | --- | --- |
| Q5NZ01 | *Aromatoleum aromaticum* EbN1 | 653 |
| Q3SJ28 | *Thiobacillus denitrificans* ATCC 25259 | 646 |
| A0A0S2SW80 | *Rhodoferax ferrireducens* | 161 |
| A0A096YHD1 | *Burkholderia thailandensis* ATCC 700388 | 654 |
| A0A0H2WK22 | *Burkholderia mallei* ATCC 23344 | 660 |
| Q63UJ3 | *Burkholderia pseudomallei* K96243 | 655 |
| A3N9S4 | *Burkholderia pseudomallei* 668 | 693 |
| A3NVK9 | *Burkholderia pseudomallei* 1106a | 655 |
| Q3JS03 | *Burkholderia pseudomallei* 1710b | 691 |
| C4KLW8 | *Burkholderia pseudomallei* MSHR346 | 655 |
| A0A0C3HZ83 | *Thauera* sp. SWB20 | 652 |
| E1VFI9 | *gamma proteobacterium* HdN1 | 654 |
| A1KA74 | *Azoarcus* sp. BH72 | 644 |
| A1W547 | *Acidovorax* sp. JS42 | 646 |
| B9MFH4 | *Acidovorax ebreus* TPSY | 645 |
| F4GDV5 | *Alicycliphilus denitrificans* DSM 14773 / CIP 107495 / K601 | 644 |
| B1Y809 | *Leptothrix cholodnii* ATCC 51168 / LMG 8142 / SP-6 | 644 |
| Q59105 | *Cupriavidus necator* ATCC 17699 / H16 / DSM 428 / Stanier 337 | 643 |
| Q1LDJ7 | *Cupriavidus metallidurans* ATCC 43123 / DSM 2839 / NBRC 102507 / CH34 | 646 |
| B2UHW1 | *Ralstonia pickettii* 12J | 649 |
| B8GM53 | *Thioalkalivibrio sulfidiphilus* HL-EbGR7 | 628 |
| I4VZB7 | *Rhodanobacter fulvus* Jip2 | 647 |
| A0A154QKM8 | *Rhodanobacter thiooxydans* | 647 |
| I4W3I5 | *Rhodanobacter spathiphylli* B39 | 648 |
| D3X7S2 | *Rhodanobacter denitrificans* | 582 |
| I4VI23 | *Rhodanobacter* sp. 115 | 649 |
| A1SUT0 | *Psychromonas ingrahamii* 37 | 630 |
| Q47UZ6 | *Colwellia psychrerythraea* 34H / ATCC BAA-681 | 617 |
| Q6LJ11 | *Photobacterium profundum* SS9 | 634 |
| Q12M27 | *Shewanella denitrificans* OS217 / ATCC BAA-1090 / DSM 15013 | 628 |
| A3QIG8 | *Shewanella loihica* ATCC BAA-1088 / PV-4 | 628 |
| Q0A9R1 | *Alkalilimnicola ehrlichii* ATCC BAA-1101 / DSM 17681 / MLHE-1 | 640 |
| A1U583 | *Marinobacter hydrocarbonoclasticus* ATCC 700491 / DSM 11845 / VT8 | 631 |
| Q2SGT6 | *Hahella chejuensis* KCTC 2396 | 639 |
| Q9HYL2 | *Pseudomonas aeruginosa* ATCC 15692 / DSM 22644 / CIP 104116 / JCM 14847 / LMG 12228 / 1C / PRS 101 / PAO1 | 636 |
| A0A0H2ZE88 | *Pseudomonas aeruginosa* UCBPP-PA14 | 636 |
| A6V237 | *Pseudomonas aeruginosa* PA7) | 636 |
| Q01710 | *Pseudomonas aeruginosa* | 634 |
| A4VQC7 | *Pseudomonas stutzeri* A1501 | 637 |
| F4DW40 | *Pseudomonas mendocina* NK-01 | 634 |
| F2KF53 | *Pseudomonas brassicacearum* NFM421 | 645 |
| G8Q381 | *Pseudomonas fluorescens* F113 | 628 |
| G8PG62 | *Pseudovibrio* sp. FO-BEG1 | 609 |
| Q16A17 | *Roseobacter denitrificans* ATCC 33942 / OCh 114 | 637 |
| F2J4Q5 | *Polymorphum gilvum* LMG 25793 / CGMCC 1.9160 / SL003B-26A1 | 632 |
| A4EUR9 | *Roseobacter* sp. SK209-2-6 | 637 |
| Q5LLH7 | *Ruegeria pomeroyi* ATCC 700808 / DSM 15171 / DSS-3 | 635 |
| A8LM18 | *Dinoroseobacter shibae* DSM 16493 / NCIMB 14021 / DFL 12 | 647 |
| A4WXT8 | *Rhodobacter sphaeroides* ATCC 17025 / ATH 2.4.3 | 644 |
| B9KWZ3 | *Rhodobacter sphaeroides* KD131 / KCTC 12085 | 644 |
| D5AVJ6 | *Rhodobacter capsulatus* ATCC BAA-309 / NBRC 16581 / SB1003 | 648 |
| A1B9T9 | *Paracoccus denitrificans* Pd 1222 | 652 |
| A0A1V0DTB3 | *Neisseria lactamica* | 655 |
| E3HH49 | *Achromobacter xylosoxidans* A8 | 638 |
| A9ICP1 | *Bordetella petrii* ATCC BAA-461 / DSM 12804 / CCUG 43448 | 635 |
| B6IY80 | *Rhodospirillum centenum* ATCC 51521 / SW | 642 |
| A0A060DX30 | *Azospirillum brasilense* | 653 |
| G7ZD37 | *Azospirillum lipoferum* 4B | 649 |
| A5EP51 | *Bradyrhizobium* sp. BTAi1 / ATCC BAA-1182 | 639 |
| Q59746 | *Rhizobium meliloti* 1021 | 639 |
| A6X742 | *Ochrobactrum anthropi* ATCC 49188 / DSM 6882 / JCM 21032 / NBRC 15819 / NCTC 12168 | 638 |
| A9ME28 | *Brucella canis* ATCC 23365 / NCTC 10854 | 639 |
| C7LHB8 | *Brucella microti* CCM 4915 | 639 |
| A0A0H3AUQ1 | *Brucella ovis* ATCC 25840 / 63/290 / NCTC 10512 | 639 |
| Q8FX16 | *Brucella suis biovar 1* 1330 | 639 |
| B0UIC2 | *Methylobacterium* sp. 4-46 | 645 |
| D8JZ83 | *Hyphomicrobium denitrificans* ATCC 51888 / DSM 1869 / NCIB 11706 / TK 0415 | 641 |
| B6JBU9 | *Oligotropha carboxidovorans* ATCC 49405 / DSM 1227 / KCTC 32145 / OM5 | 647 |
| Q89XJ6 | *Bradyrhizobium diazoefficiens* JCM 10833 / IAM 13628 / NBRC 14792 / USDA 110 | 650 |
| Q07M02 | *Rhodopseudomonas palustris* BisA53 | 648 |
| E6VCU4 | *Rhodopseudomonas palustris* DX-1 | 645 |
| B3QDW5 | *Rhodopseudomonas palustris* TIE-1 | 645 |
| Q21C83 | *Rhodopseudomonas palustris* BisB18 | 649 |

Table S5. Putative active N_2_O-reducers identified in 16S rRNA gene amplicon sequencing of the enrichments incubated with continuous streams of N_2_ gas carrying 20 ppm, 200 ppm, or 10,000 ppm N_2_O.

| **N_2_O concentration (ppmv)** | **Phylum** | **Class** | **Order** | | **Family** | **Genus** | ***nosZ* clade** | **Relative**  **abundance (%)** |
| --- | --- | --- | --- | --- | --- | --- | --- | --- |
| 20 | Proteobacteria | Alphaproteobacteria | | Rhodobacterales | *Rhodobacteraceae* | *Rhodobacter* | I | 1.9 |
| 20 | Proteobacteria | Betaproteobacteria | | Burkholderiales | *Comamonadaceae* | *Acidovorax* | I | 13.5 |
| 20 | Bacteroidetes | Flavobacteriia | | Flavobacteriales | *Flavobacteriaceae* | *Cloacibacterium* | II | 18.9 |
| 20 | Bacteroidetes | Flavobacteriia | | Flavobacteriales | *Flavobacteriaceae* | *Flavobacterium* | II | 14.2 |
| 20 | Ignavibacteriae | Ignavibacteria | | Ignavibacteriales | *Ignavibacteriaceae* | *Ignavibacterium* | II | 0.6 |
| 20 | Proteobacteria | Betaproteobacteria | | Rhodocyclales | *Rhodocyclaceae* | *Azospira* | II | 2.6 |
| 200 | Proteobacteria | Gammaproteobacteria | | Pseudomonadales | *Pseudomonadaceae* | *Pseudomonas* | I | 0.4 |
| 200 | Proteobacteria | Gammaproteobacteria | | Pseudomonadales | *Moraxellaceae* | *Acinetobacter* | I | 5.6 |
| 200 | Proteobacteria | Betaproteobacteria | | Burkholderiales | *Oxalobacteraceae* | *Massilia* | I | 0.5 |
| 200 | Proteobacteria | Betaproteobacteria | | Burkholderiales | *Comamonadaceae* | *Diaphorobacter* | I | 1.9 |
| 200 | Bacteroidetes | Flavobacteriia | | Flavobacteriales | *Flavobacteriaceae* | *Flavobacterium* | II | 15.8 |
| 200 | Bacteroidetes | Flavobacteriia | | Flavobacteriales | *Flavobacteriaceae* | *Cloacibacterium* | II | 6.4 |
| 200 | Proteobacteria | Betaproteobacteria | | Rhodocyclales | *Rhodocyclaceae* | *Dechloromonas* | II | 17.3 |
| 200 | Proteobacteria | Betaproteobacteria | | Rhodocyclales | *Rhodocyclaceae* | *Azospira* | II | 2.9 |
| 10,000 | Proteobacteria | Alphaproteobacteria | | Rhodobacterales | *Rhodobacteraceae* | *Rhodobacter* | I | 3.1 |
| 10,000 | Proteobacteria | Gammaproteobacteria | | Pseudomonadales | *Moraxellaceae* | *Acinetobacter* | I | 0.3 |
| 10,000 | Proteobacteria | Betaproteobacteria | | Burkholderiales | *Comamonadaceae* | *Acidovorax* | I | 9.9 |
| 10,000 | Proteobacteria | Betaproteobacteria | | Hydrogenophilales | *Hydrogenophilaceae* | *Thiobacillus* | I | 1.1 |
| 10,000 | Bacteroidetes | Flavobacteriia | | Flavobacteriales | *Flavobacteriaceae* | *Chryseobacterium* | II | 1.3 |
| 10,000 | Bacteroidetes | Flavobacteriia | | Flavobacteriales | *Flavobacteriaceae* | *Flavobacterium* | II | 0.5 |
| 10,000 | Proteobacteria | Betaproteobacteria | | Rhodocyclales | *Rhodocyclaceae* | *Dechloromonas* | II | 45.9 |
| 10,000 | Proteobacteria | Betaproteobacteria | | Rhodocyclales | *Rhodocyclaceae* | *Azospira* | II | 0.8 |
| 10,000 | Proteobacteria | Betaproteobacteria | | Rhodocyclales | *Rhodocyclaceae* | *Thauera* | I & II | 0.3 |

Table S6. The electrophoresis gel images of the amplicons obtained from PCR of the model organisms harboring NosZG1-5 *nosZ* genes

| **Groups** | NosZG1 | NosZG2 | NosZG3 | NosZG4 | NosZG5 |
| --- | --- | --- | --- | --- | --- |
| **Model**  **Organism** | *Pseudomonas stutzeri* | *Acidovorax soli* | *Flavobacterium aquatile* | *Ignavibacterium album* | *Dechloromonas aromatica* |
| **Amplicon length** | 261 bp | 254 bp | 149 bp | 255 bp | 237 bp |
| **Gel image** | 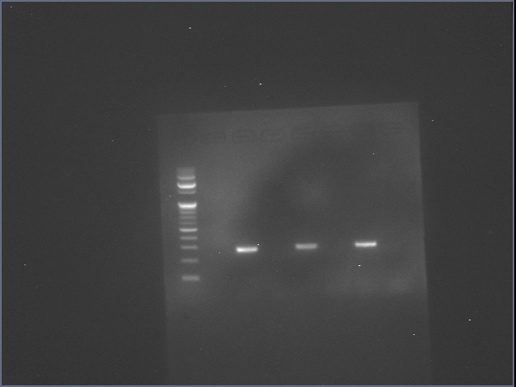 100bp  200bp  300bp  400bp  500bp | 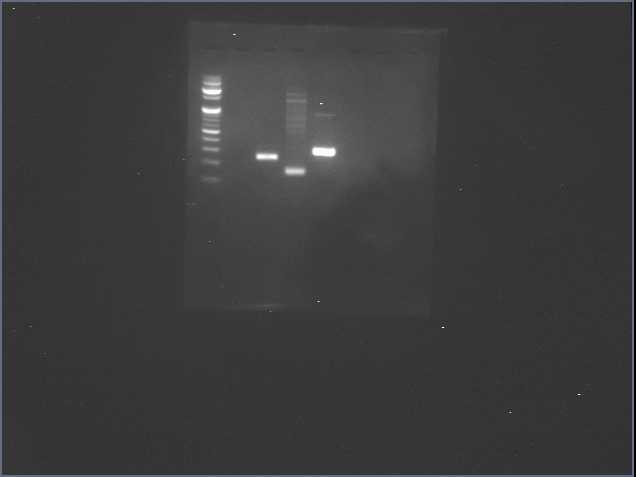 | 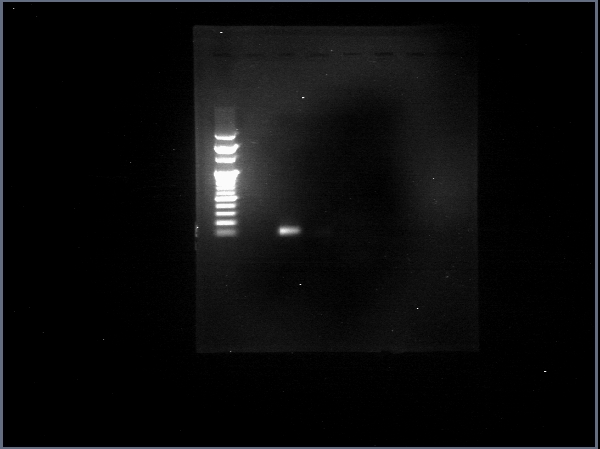 | 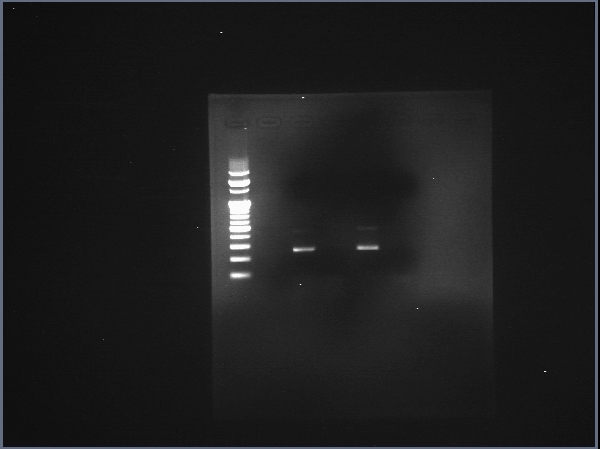 | 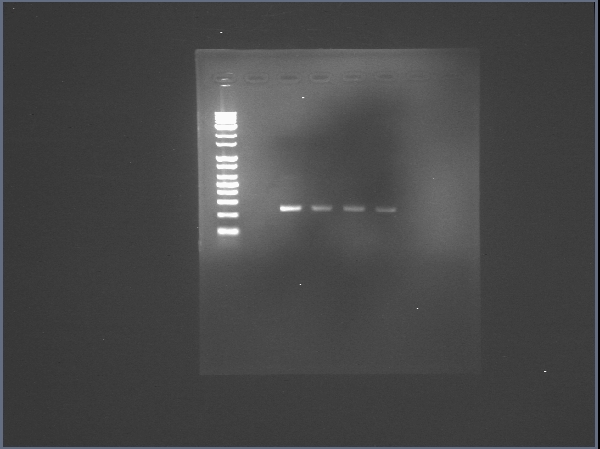 |

Table S7. Assembly statistics for the *nosZ* contigs assembled from the reads distributed to the *nosZ* bins. Only the contigs with >200bp length were considered.

| **Sample** | | **Trimmed**  **reads** | **Assembly (bp)** | | | | | |
| --- | --- | --- | --- | --- | --- | --- | --- | --- |
|  |  |  | **Sum** | **Min** | **Med** | **Mean** | **Max** | **N50** |
| DJ1 | Clade I | 276 | 105501 | 207 | 267 | 382 | 2704 | 387 |
|  | Clade II-1 | 606 | 246012 | 202 | 289 | 405 | 2624 | 412 |
|  | Clade II-2 | 635 | 267972 | 200 | 284 | 422 | 2817 | 442 |
| DJ2 | Clade I | 194 | 70179 | 207 | 270 | 361 | 2520 | 344 |
|  | Clade II-1 | 428 | 162502 | 201 | 281 | 379 | 2301 | 384 |
|  | Clade II-2 | 431 | 186144 | 207 | 290 | 431 | 2736 | 488 |
| Gwangju | Clade I | 162 | 60057 | 207 | 277 | 370 | 1281 | 359 |
|  | Clade II-1 | 321 | 131049 | 201 | 273 | 408 | 2787 | 424 |
|  | Clade II-2 | 396 | 165413 | 204 | 274 | 417 | 2925 | 428 |
| Gapyeong | Clade I | 128 | 48278 | 207 | 260 | 377 | 2333 | 358 |
|  | Clade II-1 | 260 | 100686 | 207 | 271 | 387 | 2699 | 370 |
|  | Clade II-2 | 277 | 116917 | 207 | 277 | 422 | 2759 | 462 |

Table S8. Putative active N_2_O-reducers identified in 16S rRNA gene amplicon sequencing of the soil enrichments incubated with continuous stream of N_2_ gas carrying 20 ppm N_2_O (>0.3% relative abundance)

| **Phylum** | **Class** | **Order** | **Family** | **Genus** | **Relative abundance (%)** | **NosZ**  **Group** |
| --- | --- | --- | --- | --- | --- | --- |
| Proteobacteria | Gammaproteobacteria | Pseudomonadales | *Pseudomonadaceae* | *Pseudomonas* | 59.3 | NosZG1 |
| Proteobacteria | Betaproteobacteria | Burkholderiales | *Comamonadaceae* | *Acidovorax* | 31.1 | NosZG2 |
| Proteobacteria | Betaproteobacteria | Burkholderiales | *Comamonadaceae* | *Ramlibacter* | 2.1 | NosZG2 |
| Bacteroidetes | Flavobacteriia | Flavobacteriales | *Flavobacteriaceae* | *Chryseobacterium* | 1.9 | NosZG3 |
| Proteobacteria | Betaproteobacteria | Rhodocyclales | *Rhodocyclaceae* | *Shinella* | 1.1 | NosZG1 |
| Proteobacteria | Alphaproteobacteria | Rhizobiales | *Rhizobiaceae* | *Rhizobium* | 0.7 | NosZG1 |
| Bacteroidetes | Flavobacteriia | Flavobacteriales | *Flavobacteriaceae* | *Flavobacterium* | 0.3 | NosZG3 |
| Proteobacteria | Betaproteobacteria | Burkholderiales | *Oxalobacteraceae* | *Janthinobacterium* | 0.3 | Putative NosZG2 |

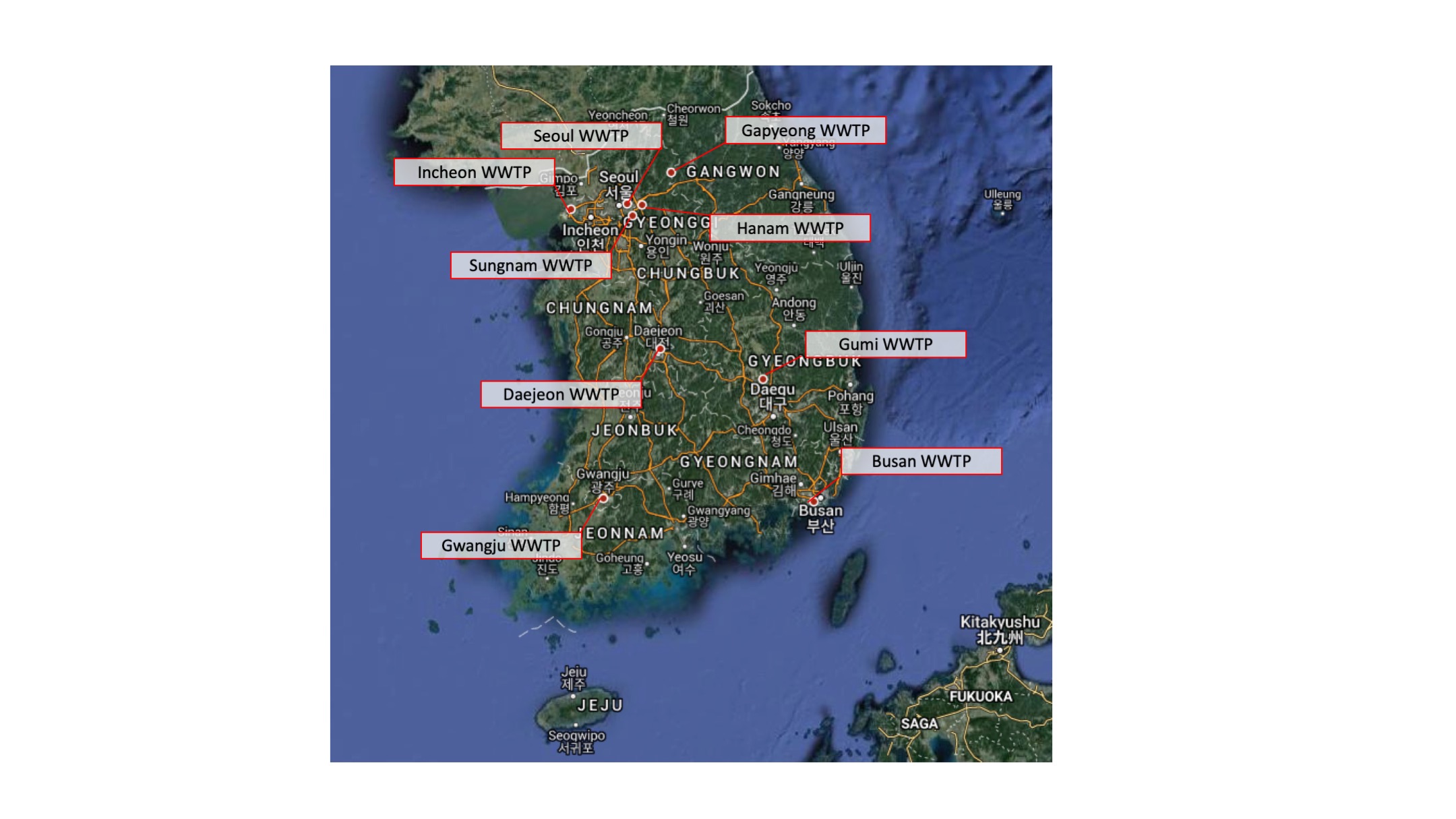

Figure S1. The location of the nine A2O wastewater treatment plants in Korea, where the activated sludge samples for *nosZ* quantification were collected.

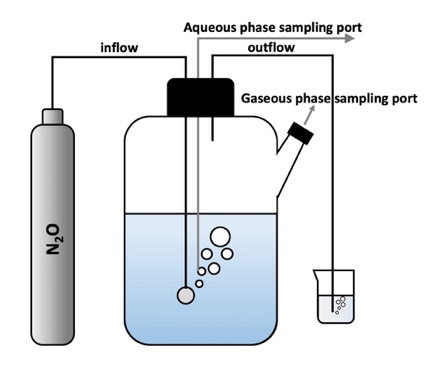

Figure S2. The fed-batch bioreactor used for enrichment of N_2_O-reducing organisms in Daejeon1 activated sludge sample. The reactor was fed with continuous stream of N_2_O-carrying gas prepared with N_2_ as the background gas.

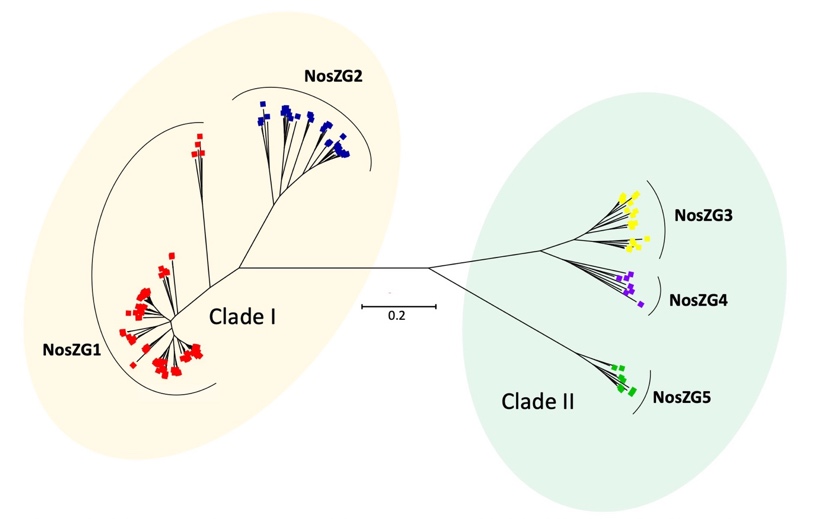

Figure S3**.** The phylogenetic tree of 174 *nosZ* sequences belonging to 14 genera enriched after fed-batch incubation with N_2_O. The *nosZ* sequences were imported from full, draft, and metagenome-assembled genome sequences imported from the NCBI database and the tree was built using neighbor-joining algorithm with 500 bootstrap replicates

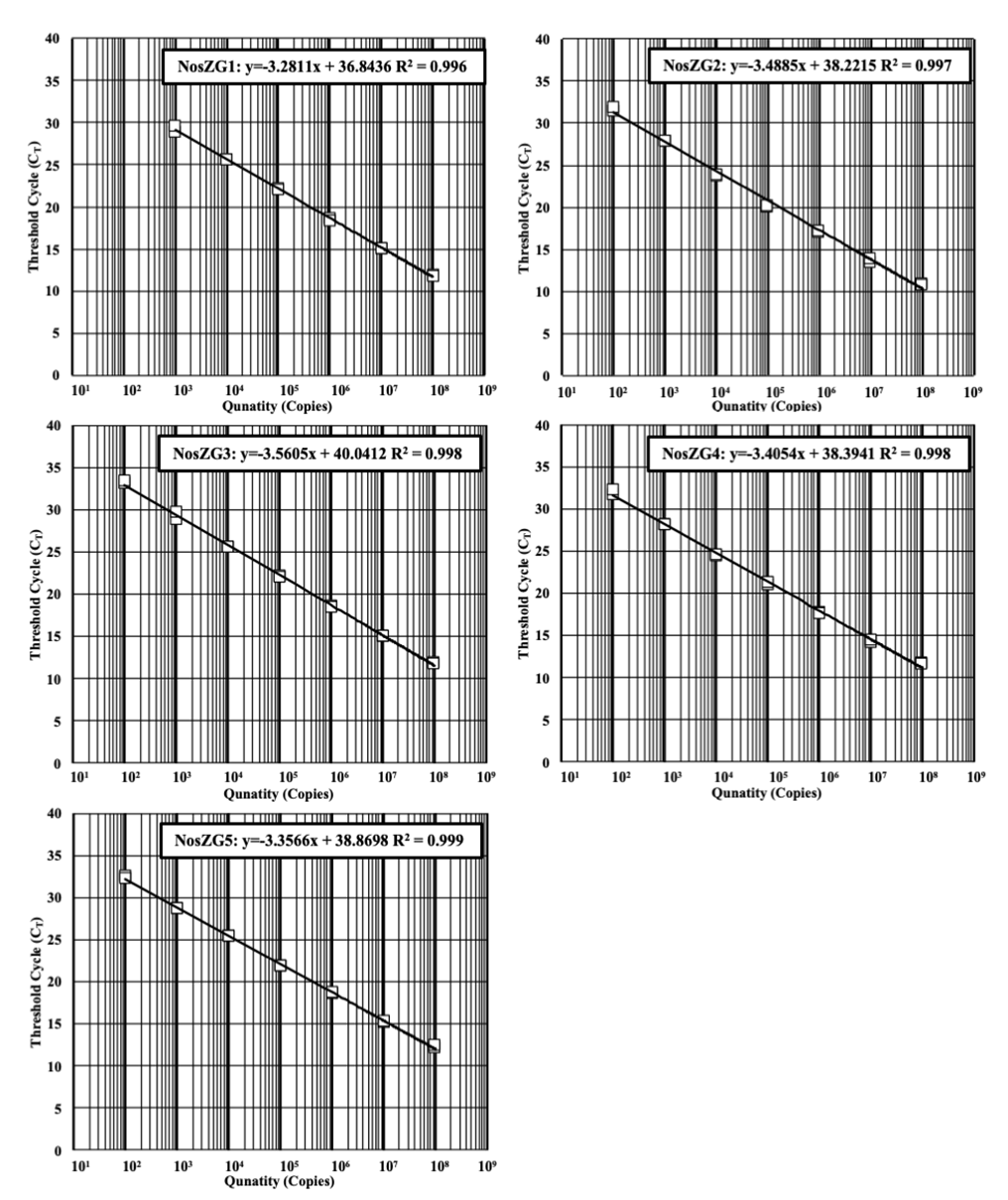

Figure S4. The calibration curves for TaqMan qPCR targeting *nosZ* genes of the model organisms (NosZG1: *Pseudomonas stutzeri* DCP-Ps1; NosZG2: *Acidovorax soli* DSM25157; NosZG3: *Flavobacterium aquatile* LMG4008; NosZG4: *Ignavibacterium album* JCM16511; NosZG5: *Dechloromonas aromatica* RCB) with NosZG1-5 primer/probe sets

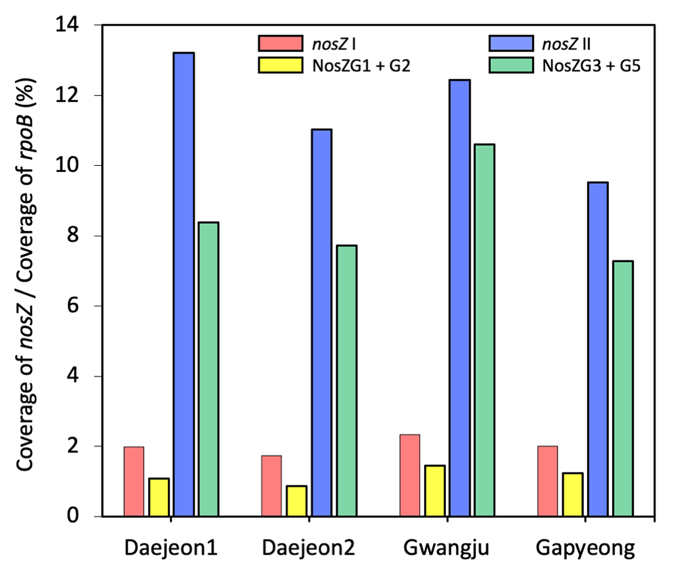

Figure S5. The relative abundances of clade I and clade II *nosZ* sequences extracted from the metagenomes of the four activated sludge samples. The bars labeled as *nosZ* I (Red) and *nosZ* II (Blue) represent the total abundances of the sequences binned as clade I and clade II *nosZ*, respectively. The bars labeled as NosZG1+G2 (Yellow) and NosZG3+G5 (Green) represent the abundances of the contigs sharing the same annotations with the OTUs generated from sequencing of NosZG1/2 and NosZG3/5 amplicons, respectively

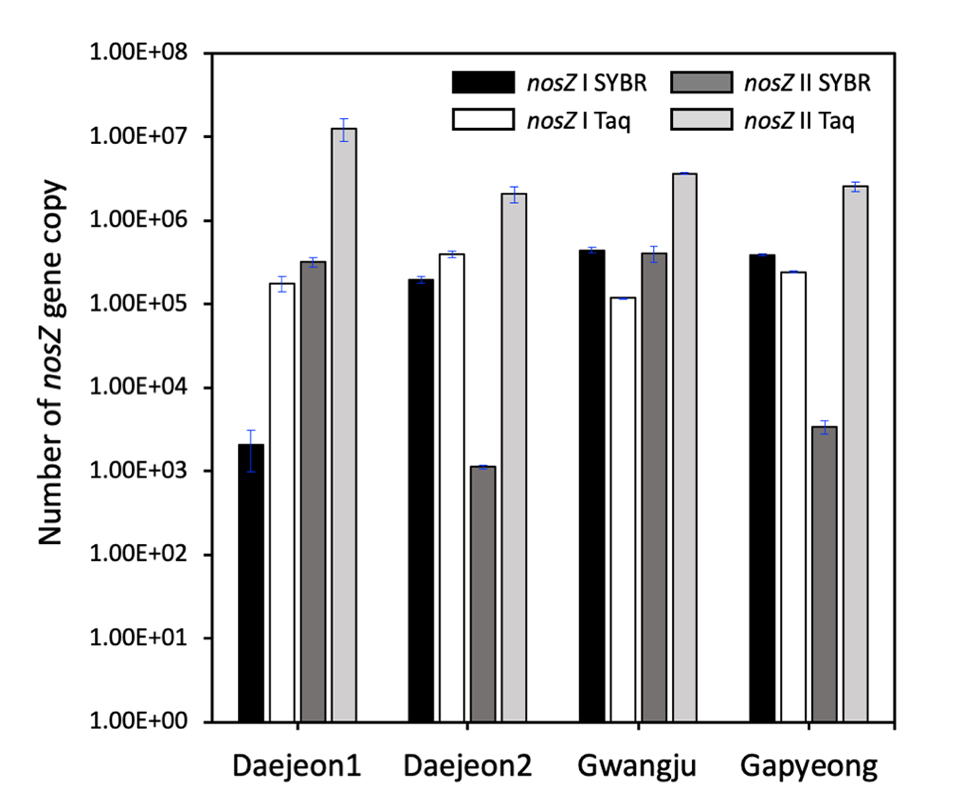

Figure S6. Copy numbers of clade I and clade II *nosZ* genes, as quantified with the SYBR Green qPCR using 1840F/2090R (clade I) and nosZII-F/nosZII-R (clade II) primer sets and TaqMan qPCR using NosZG1-5 primer/probe sets. Clade I *nosZ* gene copy numbers are sums of NosZG1 and NosZG2 qPCR results and clade II *nosZ* genes are sums of NosZG3 and NosZG4 qPCR results. The presented qPCR quantification data are the averages of triplicate samples processed seperately through extraction and qPCR procedures, with the error bars representing their standard deviations.

**References**

1. Ritalahti, K.M.; Amos, B.K.; Sung, Y.; Wu, Q.; Koenigsberg, S.S.; Löffler, F.E. Quantitative PCR targeting 16S rRNA and reductive dehalogenase genes simultaneously monitors multiple *Dehalococcoides* strains. *Appl. Environ. Microbiol.* **2006**, *72*, (4), 2765-2774.
2. Henry, S.; Bru, D.; Stres, B.; Hallet, S.; Philippot, L. Quantitative detection of the *nosZ* gene, encoding nitrous oxide reductase, and comparison of the abundances of 16S rRNA, *narG*, *nirK*, and *nosZ* genes in soils. *Appl. Environ. Microbiol.* **2006**, *72*, (8), 5181-5189.
3. Jones, C.M.; Graf, D.R.; Bru, D.; Philippot, L.; Hallin, S. The unaccounted yet abundant nitrous oxide-reducing microbial community: a potential nitrous oxide sink *ISME J.* **2013**, *7*, (2), 417.
4. Johnson, D.R.; Lee, P.K.; Holmes, V.F.; Alvarez-Cohen, L. An internal reference technique for accurately quantifying specific mRNAs by real-time PCR with application to the *tceA* reductive dehalogenase gene. *Appl. Environ. Microbiol,* **2005**, *71*, (7), 3866-3871.
